# Are drug targets with genetic support twice as likely to be approved? Revised estimates of the impact of genetic support for drug mechanisms on the probability of drug approval

**DOI:** 10.1101/513945

**Authors:** Emily A. King, J. Wade Davis, Jacob F. Degner

**Affiliations:** AbbVie Genomics Research Center, North Chicago, IL

## Abstract

Despite strong vetting for disease activity, only 10% of candidate new molecular entities in early stage clinical trials are eventually approved. Analyzing historical pipeline data, Nelson et al. 2015 (Nat. Genet.) concluded pipeline drug targets with human genetic evidence of disease association are twice as likely to lead to approved drugs. Taking advantage of recent clinical development advances and rapid growth in GWAS datasets, we extend the original work using updated data, test whether genetic evidence predicts future successes and introduce statistical models adjusting for target and indication-level properties. Our work confirms drugs with genetically supported targets were more likely to be successful in Phases II and III. When causal genes are clear (Mendelian traits and GWAS associations linked to coding variants), we find the use of human genetic evidence increases approval from Phase I by greater than two-fold, and, for Mendelian associations, the positive association holds prospectively. Our findings suggest investments into genomics and genetics are likely to be beneficial to companies deploying this strategy.

## 2 Introduction

The cost of developing new molecular entities (NMEs) into approved therapies continues to sky rocket with cost per launched NME ranging from $3 billion to more than $10 billion across major research based pharmaceutical companies [26]. Despite strong vetting for disease activity, only 5-10% of candidate NMEs in early stage clinical trials are eventually approved and this probability of approval has a direct relationship to total cost per approved drug [23, 26]. Thus, to maintain a sustainable drug development process, there is a critical need to increase the number of successful NMEs, while reducing the number of failures.

Analyzing historic data of the progress of drug compounds through the drug development pipeline, Nelson et al. 2015 [21] concluded pipeline drug targets with human genetic evidence of disease association are twice as likely to lead to approved drugs. The specific claim of doubled approval probability, if true, could lead to fewer failed clinical programs thereby lowering drug development costs. Indeed, using the estimated impact of genetics from Nelson et al. [21], increasing the fraction of NMEs in development with genetic support from the current value of 15% to 50% is predicted to decrease the direct R&D cost per launched drug by 22 ± 13% [15].

Several recent successes have corroborated the power of leveraging genetic data to predict the success of a new drug targets. Genetic evidence linking mutations in the LDL receptor gene (LDLR) to high LDL cholesterol levels and increased risk of heart disease led to lovastatin and many other drugs that inhibit HMG-CoA reductase, a rate limiting step in LDL biosynthesis [30]. The gain of function mutations in PCSK9 [7, 2, 16, 8], which cause familial hypercholesterolaemia and coronary artery disease led to to the launch of Evolocumab (Amgen) and Alirocumab (Regeneron). In question is how widely the pharmaceutical industry can expect genetics and genomics to yield increased success rates beyond these more narrowly defined examples that have unambiguous causal genes and multiple verified Mendelian mutations. If the association between human genetic evidence and approved drugs is genuine and continues to hold for present-day drug development, we expect better variant to gene mapping methods and more sophisticated predictive approaches will further improve our ability to prioritize drug targets. Because of the foundational nature of the Nelson et al. work [27], it is important to determine whether the reported association holds prospectively, and whether it replicates on independent data subsets not used in the original model construction.

Three years have passed since the publication by Nelson et al. and five years have passed since the data freeze used for analysis happened [21]. The results may now be validated using drug progression events to which Nelson et al. were completely blinded at the time. Similarly, ongoing efforts in discovering disease-associated variants in increasingly large patient samples have rapidly grown the number of potential gene trait links. For example, a public central repository of genetic association studies (GWAS Catalog [19], https://www.ebi.ac.uk/gwas/) has grown by four-fold [33, 18]. Additionally, the quantity and quality of links between noncoding SNPs and genes has expanded with the development of GTEx [13]. Here we report revised estimates of the impact of genetic evidence on drug target success and extend Nelson’s observations into a model that can be deployed by other companies and academics to predict the likelihood of success of targets of interest to them.

## 3 Results

### 3.1 Identifying Validation Sets

Nelson et al. [21] estimated a twofold increase in approval probability for Phase I drug targets with genetic evidence using drug pipeline data from Informa Pharmaprojects along with genetic data from a variety of sources, all obtained in 2013. This estimate comes from historical rather than experimental data so a direct replication is not possible. However, we can obtain updated sources of pipeline and genetic data and use the data subsets not used in the Nelson et al. study study to validate its claims. Figure 1A shows how updated pipeline (Informa Pharmaprojects [1]) and genetic association (GWAS Catalog, OMIM [20]) datasets may be split into discrete subsets, several of which were not used in the original analysis. We call these sets validation sets. In addition to genetic associations and pipeline progression events added after 2013 (New Genetic and Pipeline Progression sets), we identified a large subset of pipeline data that was available to Nelson et al., but that was excluded from analysis because Pharmaprojects reported an inactive status, most commonly “No Development Reported”. Instead of directly using Pharmaprojects development status, we use other fields in the database to label drugs with a latest historical development phase (see Methods, S2.1), enabling us to use 83% of this data in our analysis.

**Figure 1:**
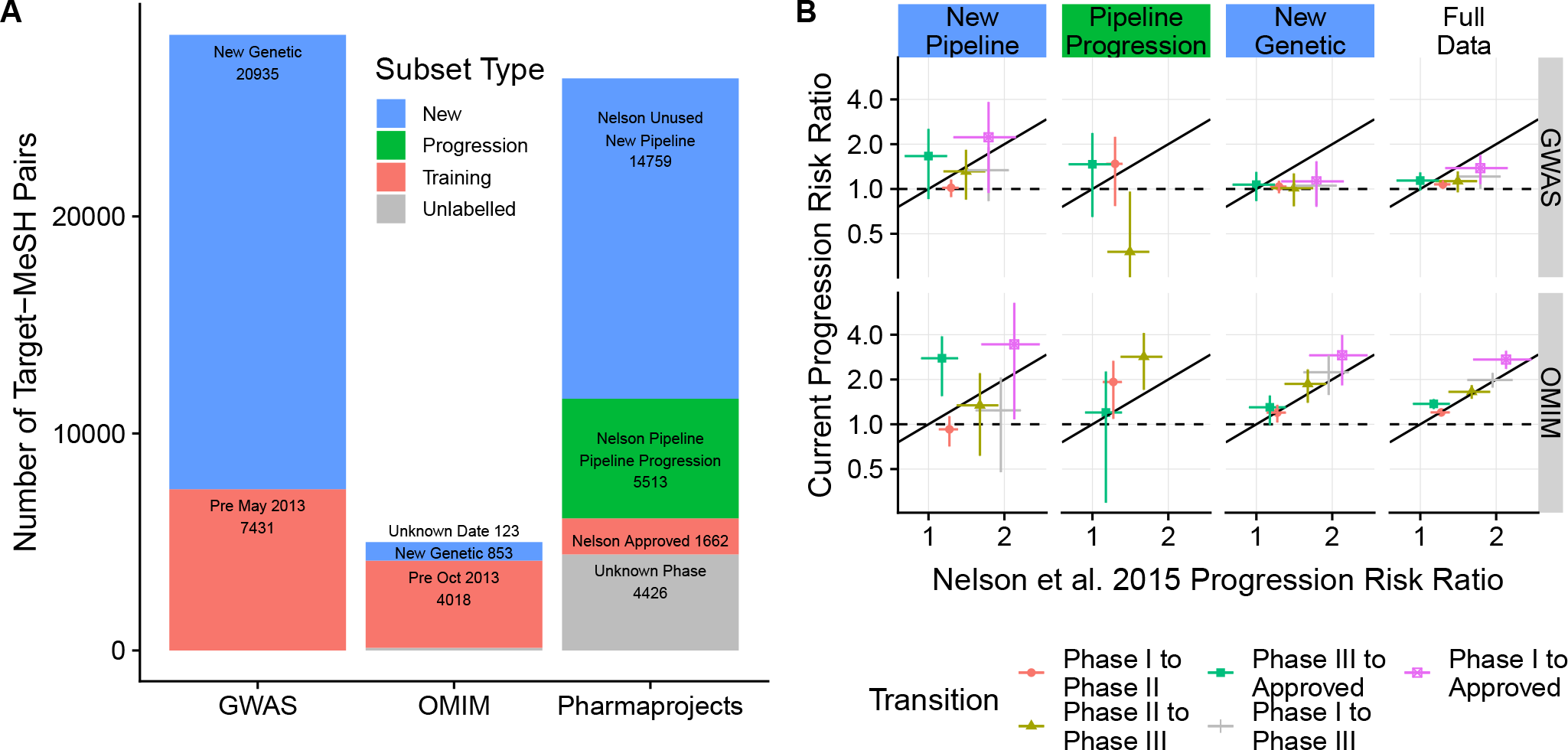
Estimated effect of evidence from human genetic studies on the probability of advancing in clinical development. A: Partitioning Pharmaprojects, OMIM, and GWAS Catalog into training data available to Nelson et al. 2015 and validation sets. We use validation set Pipeline Progression (which drugs advanced >2013) to determine whether gene target-indication pairs with genetic evidence were more likely to advance to the next pipeline phase from 2013-2018. B: Our estimates of the effect of genetic evidence on gene target-indication pair progression compared to values reported by Nelson et al. 2015 [21] in validation sets New Pipeline (drugs and indications > 2013, 2013 inactive drugs) New Genetic (only new genetic information > 2013) Pipeline Progression, and in the full updated dataset (Full Data).

Following Nelson, we aggregate data at the level of gene target-indication pair, the unit on which genetic evidence is computed. In total, we mapped 26360 gene target-indication pairs to a highest pipeline phase, in contrast to 8853 pairs labelled with an known phase in the Nelson et al. analysis. 5513 pairs could be tested for progression to a more advanced clinical phase since 2013, and 14759 pairs either absent or inactive in the 2013 data set could now be assigned a highest historical pipeline phase. Two validation sets (New Pipeline, and new GWAS associations) are larger than the original datasets used in Nelson et al. giving us sufficient power to test predictions.

Our replication analysis occurred in three steps. In the first step, we took labels of genetic evidence directly from Nelson et al. 2015 and tested how these labels predict pipeline outcomes in the New Pipeline and Pipeline Progression validation sets. Second, we repeated the analysis using both updated pipeline data and updated genetic association datasets and determined whether genetic evidence labels constructed from associations reported after 2013 are positively associated with historical progression. This analysis uses the New Genetic validation set, defined as GWAS data added after May 2013 and OMIM data added after October 2013. Third, we determine whether genetic labels constructed from the full set of updated GWAS and OMIM genetic associations are linked to improved pipeline outcomes over the entire updated Pharmaprojects dataset (See Methods and Supplemental Data for more details). We refer to this analysis as Full Data.

### 3.2 Estimated Effect of Genetic Evidence on Validation sets

Of the many results from the original Nelson et al. publication, we focus on determining whether the probability of progressing along the development pipeline is greater for gene target-indication pairs with genetic evidence as this most directly impacts business decision-making (supplementary figures S8-9 and S11-12 show replication of other results). A gene target-indication pair is said to have *genetic evidence* if there is human genetic evidence of association between the gene target and a trait sufficiently similar to the indication (see sections 5.3 and 5.4). Figure 1B shows estimates and 95% confidence intervals for the ratio of the probability of progression for gene target-indication pairs with and without genetic evidence computed on the three validation sets and the full set of new data each plotted against values computed from Nelson et al. supplementary tables.

Across all three validation sets (Pipeline Progression, New Genetic, and New Pipeline), we consistently see a marked difference between the effect of genetic evidence derived from the OMIM database and genetic evidence derived from the GWAS Catalog. Estimated effects of OMIM genetic evidence are comparable to or greater than previously reported values [21], except for progressions from Phase I to Phase II, which are lower using new data. Notably, we see a positive and significant effect of OMIM genetic evidence on the probability of progression from Phase II to Phase III since 2013 (Pipeline Progression validation set). With the exception of progressing from Phase III to Approval, estimated effects from GWAS Catalog-derived genetic evidence are consistently lower than the originally reported values. Our estimated effects of GWAS genetic evidence in the New Genetic validation set are often significantly lower than the originally reported values. In validation sets, all estimates of the effect of GWAS evidence overlap one (no effect), except in the Pipeline Progression validation set, where we estimate a negative effect of GWAS evidence on Phase II to III progression (Fig 1B).

In both GWAS and OMIM datasets, our estimates of the effect of genetic evidence on Phase I to II progression probabilities are lower than originally reported, and confidence intervals sometimes exclude original estimates. With some exceptions (e.g. oncology studies), Phase I trials assess safety in healthy volunteers, not efficacy, so their success may be less closely linked to human genetic evidence for target involvement in disease. Validation sets may also differ systematically from the 2013 training data. For example, it is possible that there are systematic differences in the types of associations discovered before and after 2013 (New Genetic validation set). Later associations may be biased towards those with smaller effect sizes or rarer variants only detectable in larger cohorts, and could also be less predictive of drug efficacy. Using the complete updated dataset (Full Data), including all Pharmaprojects drugs and pre and post 2013 genetic associations, we find the estimated effect of GWAS genetic evidence on Phase I to Approval is still significantly positive, and the effect of OMIM genetic evidence is greater than originally reported. With more data accumulation and a modified strategy for classifying data, we more accurately predict the likelihood of approval for gene target-indication pairs with genetic evidence.

### 3.3 Statistical modeling of genetic effect on drug approval

The effect of GWAS genetic evidence on approval was considerably reduced and lacked statistical significance in the New Genetic dataset. In reanalyzing the original data, we found the estimated effect of GWAS genetic evidence was highly sensitive to the choice of trait-indication similarity cutoff used to determine whether or not a drug target had a genetic association (Figure S3). Learning from this analysis, we sought to build a model relating genetic evidence to the probability of drug approval in the full dataset.

We fit multivariate logistic regression models predicting target-indication pair approval using several independent variables. The first was a measure of (continuous) genetic evidence, defined as the greatest similarity to the indication across all traits linked to the drug target through human genetic evidence. The remaining independent variables are target and indication-level properties that could confound the relationship between genetic evidence and approval. Previous work has shown that approved drug targets tend to be more conserved than genes linked to GWAS associations [4], so we included residual variant intolerance score (RVIS) [24], measuring the amount of common functional variation in each gene relative to the amount of neutral variation, as a predictor. We also included the amount of time each target is known to have been under development as a predictor, with the rationale that if accumulating genetic evidence informs drug development, targets supported by genetic evidence might be newer on average. Finally, we included gene ontology (GO) terms and high level MeSH terms for each indication as predictors to control for known differences [14, 28] in approval probability among indication and target classes.

Under this model, approval is positively associated with trait similarity for supporting GWAS and OMIM associations, with 95% credible intervals excluding zero (Figure 2). When associated traits are sufficiently similar, gene target-indication pairs with GWAS or OMIM associations are more likely to be approved. Evaluation of the data also revealed when there is a genetic association for a dissimilar disease, they are less likely to be approved than gene target-indication pairs with no known genetic association. This negative association is a novel finding.

**Figure 2:**
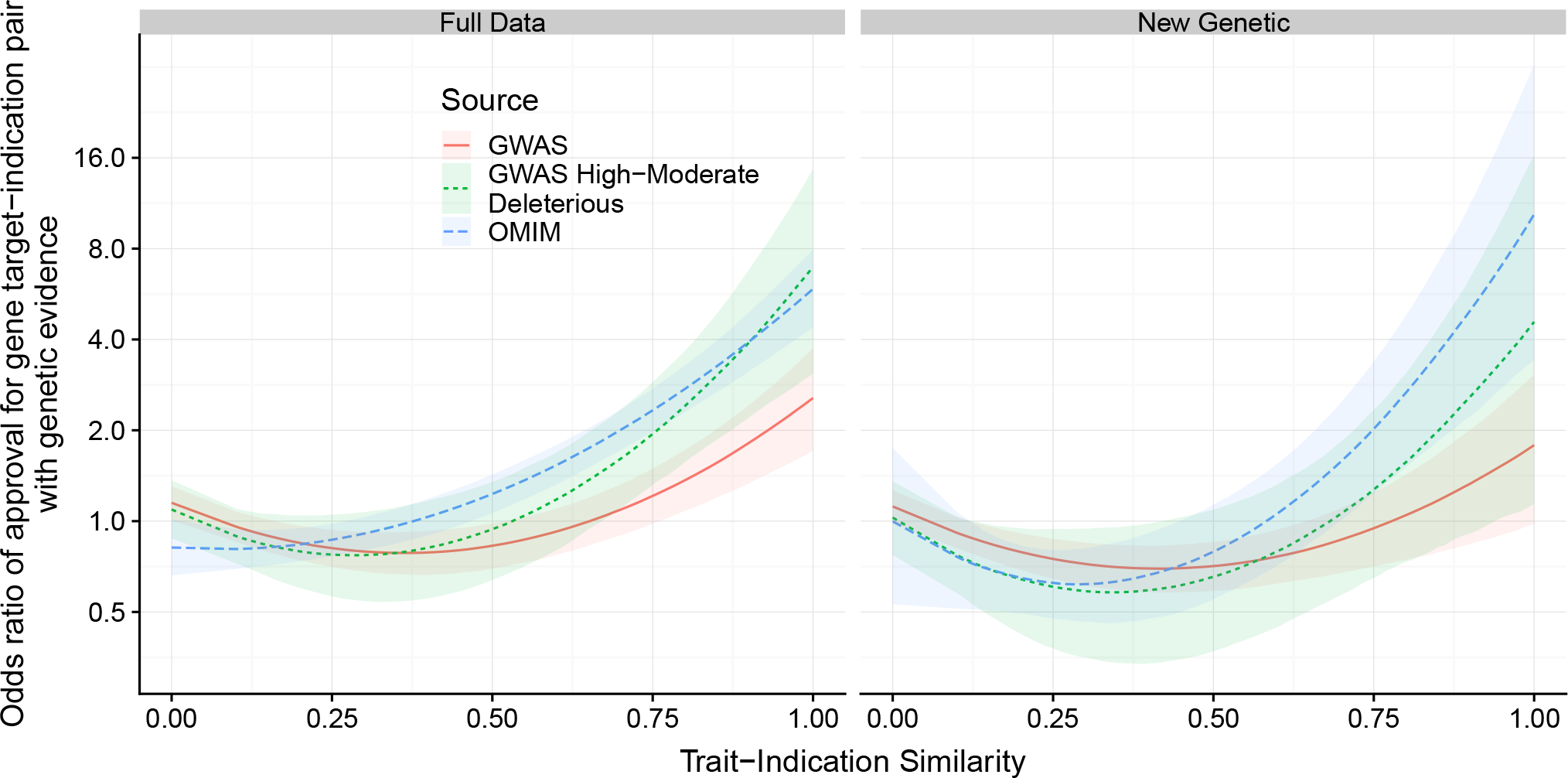
Estimated odds ratio of gene target-indication pair attaining approval, as a function of similarity between drug indication and the most similar trait associated with the target. Left: All genetic associations. Right: Only genetic associations reported after 2013 download. Posterior median and pointwise 95% credible interval from Bayesian logistic regression.

GWAS genetic evidence has a smaller positive effect on approval than does OMIM genetic evidence, and we only find a small beneficial effect of GWAS genetic evidence in the New Genetic validation set. One possible explanation is that most GWAS associations are to noncoding variants, and determining function from these associations will require more advanced methodology [11]. Indeed, when we only consider GWAS Catalog SNPs in high LD (*R*^2^ ≥ 0.9) to a missense variant or other variant predicted to be moderately or highly deleterious [6], the estimated effect of GWAS genetic evidence on drug target approval approaches that of OMIM. Moreover, for missense variants, we see a larger estimated effect of genetic evidence when using a more stringent LD cutoff to the lead SNP (Figure S24).

## 4 Discussion

Pharmaceutical companies are investing in the creation and analysis of genomics data in the hope of improving target selection and decreasing failures due to lack of efficacy [10] or adverse effects [22]. Previous work by Nelson et al. 2015 [21] supported this investment, showing gene target-indication pairs with genetic evidence are approximately twice as likely to progress from Phase I to approval. This quantitative estimate is the product of many decisions, for example how to identify similar traits in genomics and pipeline databases, that, although reasonable, could have been made differently. Additionally, the results were based on a large historical set of approved drugs and might not hold for present-day target selection. This motivated us to replicate the analysis using using 5 years of data that has accumulated since their data freeze in 2013.

In the replication study, we recovered a robust association between OMIM genetic evidence and drug approval of a similar or greater magnitude to that originally reported [21] across several independent test sets. GWAS genetic evidence also is generally positively associated with progressing in clinical development, but the magnitude of the association is smaller and not clearly different from zero in any independent replication set. There appears to be some confounding due to GWAS genes having different properties than successful drug targets. When this is controlled for using logistic regression, GWAS-supported target-indication pairs are more likely to succeed than those without a GWAS-linked gene target. This highlights the need for predictive models including target properties, work that is beginning to emerge [34].

The OMIM database provides expert-curated gene-trait links, bypassing the need to assign noncoding SNPs to genes, a major source of uncertainty for present GWAS methods. Better methods for linking GWAS SNPs to causal genes may improve performance, supported by the fact that we found strong and statistically significant positive associations between GWAS genetic evidence and drug success when considering only the highest confidence SNP-gene links, characterized as having a leading SNP with *R*^2^ ≥ 0.9 to a variant predicted to be highly or moderately deleterious. However, OMIM’s focus on Mendelian phenotypes also means genetic variants will be higher effect size than those for quantitative traits or conditions prominent in the GWAS Catalog, which is unlikely to be addressed by improved computational methods.

Because OMIM is a manually curated database, it is possible that known drug mechanisms influence OMIM entries, creating a positive association between OMIM genetic evidence and approval. However, we observe a positive effect of OMIM genetic associations reported by Nelson et al. 2015 on progression events occurring after data were collected for that paper, which is inconsistent with this reverse causal hypothesis. It is also possible these progression events are not truly independent of pre-2013 approvals, because they may represent approval for an indication similar to the original indication. However, the positive effect of OMIM genetic evidence on 2013-18 progression remains significant when targets with pre-2013 approvals for similar indications are excluded (Tables S11-12). We conclude the predictive effect of OMIM genetic evidence is not a statistical artifact, and is more likely to reflect the value of well-defined disease biology to drug development.

Due to the MeSH ontology structure, current methods require manual similarity assignments to recognize relationships between most quantitative traits and diseases. The high sensitivity of key results to MeSH similarity motivates treating similarity as a continuous variable and suggests improvements to its quantification. While expert curation can be advantageous in identifying closely related traits, it also leaves more room for human input to bias the analysis outcome. To assess this we removed automatically assigned similarities. Positive associations between GWAS genetic evidence and approval remain, though in some cases are greatly reduced in magnitude (Figure S16, S24) (OMIM is minimally impacted as it contains few quantitative traits). We expect improved methods automatically identifying similar phenotypes to drug indications will expand our ability to use genomics data in predictive models.

Our results highlight the importance of similarity between associated trait and drug indication in determining which gene target-indication pairs are likely to succeed. Our finding that genetic associations for highly dissimilar traits reduce the probability of approval is new and could be of significance once the reason is better understood. A possible explanation is an increased incidence of side effects due to involvement in unrelated disease mechanisms. It suggests that when target disease links are known, genetic data can improve the drug development process through improved indication selection.

Our analysis of the last five years of drug development data validates the results of Nelson et al. and indicates that the positive association between genetic evidence and drug success is not just a historical phenomenon. Using logistic regression to control for target and indication level properties, and quantifying genetic evidence on a continuous scale, we also demonstrated that associations to disparate phenotypes is a negative predictor of approval. With these algorithmic developments, we have built a Shiny [5] app that others can use to evaluate their target-indication pairs of interest. As the knowledge of what genes do biologically increases, our data suggests the reliability of genetic predictions will continue to improve. In closing, public and private investments into genomics for the purpose of improving the fraction of successful drug targets appears to be well warranted.

## 5 Methods

### 5.1 Pipeline data

Data on drug gene targets, indications, latest development phase, and approvals by country were collected from the Pharmaprojects database (accessed January 25, 2018). For each drug, Pharmaprojects provides country-level, indication-level, and global development status. The latter is the latest development status across indications for any country. A drug was considered US/EU approved for an indication if it was approved in the US or EU and approved for that indication (so if a drug is US/EU approved for one but not all of its approved indications, we will incorrectly assign some approvals). We infer this was also the approach of Nelson et al., as they mention no source other than Pharmaprojects for drug approval data and Pharmaprojects does not provide drug-indication-country level approval data.

To calculate phase-specific progression probabilities by genetic evidence, we must assign a latest historical development phase to Pharmaprojects drug-indication pairs that are not in active development using other database fields. Country status gives the latest phase for single-indication and preclinical drugs. Other drug-indication phases are determined through assessing the presence or absence of key events and clinical details matching the trial phase and the disease name. Clinical details were only used when other sources were unavailable because this field may contain information about planned or anticipated trials. Details are provided in the supplement (Section S2.1).

Pharmaprojects gene targets were mapped from Entrez to ensembl ids. Drugs with non-human and xMHC targets were excluded (following the original analysis) as were a small number of drugs with non protein coding targets.

### 5.2 Genetic data

Genetic association data was obtained from the GWAS Catalog [19] downloaded 2017-09-26. OMIM data was downloaded from [20] on 2018-06-06. GWAS Catalog associations with reported *p*-value greater than 10^−8^, OMIM provisional associations, drug response associations, and somatic variant associations were excluded.

OMIM reports gene-trait links, but the GWAS Catalog reports SNP-trait links which must be converted to gene-trait links via SNP-gene links. Although methods for creating SNP-gene links have since advanced [11], we closely follow the approach of [21] with updated data sources to reduce our degrees of freedom for overfitting to new data and to make our new estimates of the effect of genetic evidence comparable to the original estimates. Our gene-trait mapping procedure attempts to replicate that used by Nelson et al. with updated data sources. An LD expansion of GWAS Catalog reported variants was performed using an LD threshold of 0.5 in the 1000 Genomes Phase 3 EUR super population [9]. A distance-based gene-trait association was established when an LD SNP was within 5000 b.p. of the gene in hg38 as annotated by SNPEff [6]. An eQTL-based gene-trait link was established when an LD SNP was reported associated with a gene with nominal *p*-value less than 10^−6^ in any GTEx tissue [13]. Using a cutoff of 10^−12^ makes little difference to results (Table S20). A DHS-based gene-trait link was established when an LD SNP was located in a DNAse I hypersensitivity site correlated with gene expression with one-sided permutation *p*-value 1.000 (from 1000 replicates) [27]. All linked genes were mapped to Ensembl IDs, and links to genes not annotated as protein coding by Ensembl were removed from the dataset. Additional details are available in the supplement (Sections S3.1 and S3.2).

### 5.3 Trait-indication similarities

Pharmaprojects indications and GWAS Catalog and OMIM traits were mapped to MeSH headings to link traits and indications by a common vocabulary. We mapped as many terms as possible automatically by string matching to MeSH terms and their synonyms, and the remainder were manually assigned to the most specific MeSH heading encompassing the term. The MeSH vocabulary consists of MeSH headings, which are organized in a tree structure, and supplementary concepts, which are not. We did not map to MeSH supplementary concepts as the lack of structure means we cannot compute similarities between these concepts and other terms. However, each supplementary concept is assigned one or more mapped headings, and so terms matching a supplementary concept were assigned to the mapped heading. This set of MeSH term mappings was used in the full replication with new genetic data sources.

When testing predictions from the 2013 genetic association data, it was important that MeSH headings mapped to Pharmaprojects indications be consistent with the original analysis by Nelson et al. in order to correctly identify common pairs between dataset for which progression can be tested and to ensure that our New Pipeline test set contained truly novel pairs. Nelson et al. provided mappings for many Pharmaprojects indications in a supplementary dataset. Terms without provided mappings were mapped to maximize the number of Nelson et al. gene target-indication pairs also present in our dataset, subject to the mapping being biologically justifiable. Standardized mapping increased the percent of Nelson et al gene target-indication pairs present in our dataset from 88% (using our independently mapped terms) to 98%.

Resnik [25] and Lin [17] similarities between MeSH headings were computed in `R` in the `ontologySimilarity` package [12], standardized to have a maximum value of 1 for each trait, and averaged to compute a similarity between each pair of MeSH headings (Section S4.1). Two traits are considered *similar* if the similarity is greater than or equal to a critical value. Our assigned similarities are not identical to those of Nelson et al. because of using different versions of MeSH (2009 versus 2017), but were correlated with those originally reported (*R*^2^=0.86, Figure S14). We determined a critical value of 0.73 in our analysis corresponded to the critical value 0.7 used in the original analysis, and used this to determine similar traits in our replication study. Manually assigned similarities were taken from the supplement of [21]. Manual assignment was performed because the MeSH ontology makes few connections between diseases and closely related quantitative phenotypes, for example osteoporosis and bone density.

### 5.4 Genetic Evidence

We formalize and extend the concept of genetic evidence used by Nelson et al. We first define a similarity function operating on two gene-trait pairs. Define function *S* from 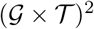 to [0, 1] where 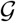 is the space of genes and 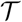 is the space of traits.

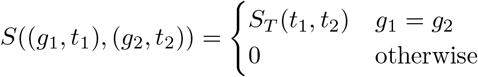

where 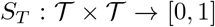 is a trait similarity function (in the Nelson et al analysis and here, computed from Resnik and Lin similarities). Let 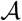 be a set of gene-trait pairs with elements in 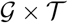 obtained from genetic data sources (for example, when analyzing the effect of OMIM genetic evidence 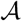 is the set of gene-trait pairs in OMIM). Genetic evidence according to Nelson et al. 2015 is a function *E*_*D*_ from 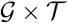 to {0, 1}

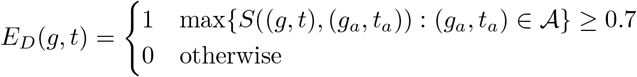

However, trait similarity is a real number in [0, 1], so we can define another genetic evidence function *E*_*C*_ from 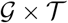 to [0, 1]

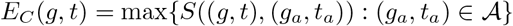

*E*_*D*_(*g*, *t*) = 1 if and only *E*_*C*_(*g*, *t*) ≥ 0.7.

### 5.5 Statistical analysis

#### 5.5.1 Two-by-two tables

Let **D** be a vector of gene target-indication-phase triplets with elements (*g*_*i*_, *t*_*i*_, *h*_*i*_), *i* = 1, …, *n*. *H*_*i*_ ∈ {0, …, 4} is an ordered categorical variable giving the latest phase each gene target-indication pair has achieved (0=Preclinical, 1=Phase I, 2=Phase II, 3=Phase III, and 4=US/EU Approved).

Risk ratios for progressing from Phase *x* to Phase *y*, *x* > *y* were computed as

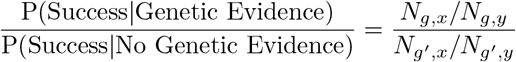

where 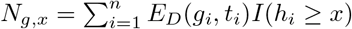 is the number of gene target-indication pairs in Phase *x* or later with genetic without genetic evidence and 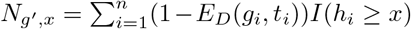 is the number of gene target-indication pairs in Phase *x* or later without genetic evidence. We required at least 5 reported genetic associations for similar traits. Phase progression probability calculations usually exclude in progress development [14] but here we include them for consistency with Nelson et al. Confidence intervals were computed using the `riskratio.boot` function in the `epitools R` package [3]. We ensured consistency of this approach with that of Nelson et al. by verifying our code could reproduce their results from supplemental materials (Section S1.1). Drugs approved only outside the US and EU and drugs with unknown latest phase were excluded from this analysis.

#### 5.5.2 Bayesian Logistic Regression

Let *i* index gene target-indication pairs (*g*_*i*_, *t*_*i*_), *i* = 1, …, *N*. Let *y*_*i*_ ∈ {0, 1} be 1 if pair *i* is found in at least one US/EU approved drug and 0 otherwise. Let *X* be an *N* × *d* design matrix where *d* is the number of non-genetic predictors with *i*^*th*^ row 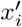.

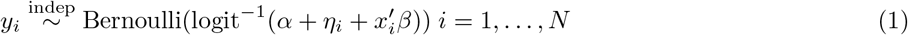

where

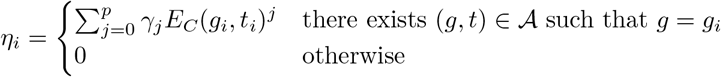

Our choice of *p* = 2 is supported by WAIC [31][32]. Predictors in **X** were top-level MeSH category, target class, estimated time the target has been under development, and RVIS score [24]. Details are provided in Section S5.1. Priors were

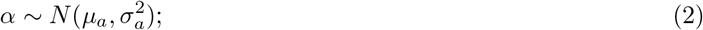

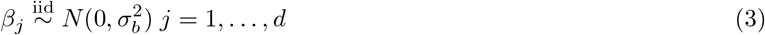

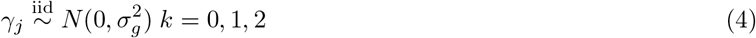

All models were fit in Stan [29] using four chains with default initialization and run settings. Prior parameters *μ*_*a*_=−2.2, *σ*_*a*_=0.75 was chosen to reflect prior knowledge that approximately 10% of Phase I compounds become approved [14] and prior standard deviations *σ*_*b*_=2, and *σ*_*g*_=2 were chosen prior belief that observed effect sizes should be moderate. Note *α*, for which we have chosen a nonzero mean prior, controls the baseline approval probability, not the effect of genetic evidence. Continuous covariates in **X** were standardized to have mean 0 and standard deviation 1 as was *E*_*C*_.

In this analysis we depart from the original Nelson et al. approach and exclude all drugs assigned an active development phase by Pharmaprojects, as it is unknown whether these development programs will ultimately lead to approval. This decision is consistent with other work estimating clinical success probabilities [14][34]. We include unapproved drugs with unknown latest historical phase. A total of 20292 gene target-indication pairs were associated with at least one US/EU approved or inactive drug and included in the analysis.

## Supporting information

Supplementary Methods And Results

## 6 Code availability

Code to reproduce the main text figures from supplementary data is provided on Github (https://github.com/AbbVie-ComputationalGenomics/genetic-evidence-approval). The git repository also contains instructions for running a Shiny app displaying model predictions.

## 7 Data availability

Supplementary data tables required to reproduce the main text figures are provided on Github (https://github.com/AbbVie-ComputationalGenomics/genetic-evidence-approval).

## 8 Acknowledgments

We thank Howard Jacob for discussions and helpful comments on the manuscript. We thank Rishi Gupta for assistance with Pharmaprojects data access.

## 9 Author contributions

E.A.K. led data acquisition and statistical modeling, wrote all computer code to process and analyze data, and developed the shiny app. J.F.D and J.W.D. conceived of the project and supervised execution. E.A.K. drafted the manuscript with support from J.F.D. and J.W.D. All authors contributed to planning the research and editing the manuscript.

