## Supplementary Methods And Results for "Are drug targets with genetic support twice as likely to be approved? Revised estimates of the impact of genetic support for drug mechanisms on the probability of drug approval"

Supplementary materials are organized as follows. In the first section, we present a replication of Nelson et al. 2015 figures from supplementary tables, and assess sensitivity to two parameters. In the second section, we describe the collection of updated pipeline data and present additional results on how 2013 genetic labels relate to clinical development outcomes (analyses New Pipeline and Pipeline Progression in the main text). In the third section, we describe collection of updated genetic data and provide additional analysis results using both updated genetic data and updated pipeline data (analyses New Genetic and Full Data in the main text). The fourth section details trait similarity methodology, and its effect on results. The final section consists of statistical modeling work. These last two analyses are performed using the full updated datasets.

### S1 Reanalysis of Nelson et al. Supplementary Datasets

#### S1.1 Replication of main text figures

| | No Genetic Association | Genetic Association |
|---|---|---|
| Not Approved | $n_{nn} = 22012 - n_{assoc} - n_{approved} + n_{aa}$ | $n_{an} = n_{assoc.} - n_{aa}$ |
| Approved | $n_{na} = n_{approved} - n_{aa}$ | $n_{aa}$ |

Table S1: Schematic of two-by-two table used in odds ratio calculation. $n_{aa}$ is directly computed as the number of genes that are both the targets of approved drugs and have reported trait associations.

**Figure 1N** In the supplementary and main results of this paper, we recreate figures from the original Nelson et al. publication with updated data sources. Before doing this, we determined if we could replicate the figures from supplementary data from the original paper (supplementary datasets 2, 3, and 4 of [20]). Figures and tables from the original publication will be referred to with the suffix N, i.e. Figure 1N. Figure 1N [20] gives the total number of MeSH, genes, and gene-MeSH pairs in the Pharmaprojects database and the GWASdb separated by source. We exactly reproduce Figure 1N from supplementary tables if sources "dbGaP", "GWAS:A", "GWAS:B", "GWASCentral", "JohnsonOdonnell" and "Omim" are considered part of the GWASdb and source "OMIM" is the only source of OMIM associations. Source "Omim" appears to have been derived from the OMIM database, although it features SNP-trait links and is largely non-overlapping with reported "OMIM" associations. We elected to exclude this data source from both the GWASdb and OMIM datasets as we wished to have a clear separation between Mendelian genetic evidence and genetic evidence from GWAS.

**Figure 2N** Figure 2N shows enrichment of approved targets among genes with known human genetic associations. Odds ratios are computed from the $2 \times 2$ table as shown in Table S1. The upper panel shows the odds ratio computed with respect to a population of 22,012 protein-coding genes, and the lower panel shows the same calculation with respect to the population of druggable genes, which we obtained from the drug-gene interaction database [10, 29]. RVIS scores for each gene were downloaded from the supplemental material of [22].

**Figure 3N and Table 1N** Figure 3N shows the proportion of gene target-indication pairs with genetic evidence by phase and by indication. Table 1N shows the risk ratio of pipeline progression for drugs with human genetic evidence. In creating these figures, Nelson et al. only included indications with at least 5 genetic associations for similar traits. The set of such indications can be computed in two ways from supplementary materials. The first approach is to compute it from supplementary data (supplementary datasets 2-4). The second approach is to refer to supplementary table 5, which gives the number of similar genetically associated trait MeSH headings for each of 704 of 705 drug indication MeSH headings (Sjogren's syndrome is missing). These two approaches yield the same number of associations per MeSH term if we define the number of genetic associations to be the number of unique Link-MSH-snp_id triplets (for OMIM, snp_id, the reported SNP, is always empty so this reduces to the number of unique Link-MSH pairs). For GWAS associations, link is a PubMed id, and for OMIM associations, link is an OMIM id. We can reproduce the results of Figure 3bN and Table 1N to within reported precision using the list of traits with at least 5 genetic associations from Supplementary Table 5 (the same as the list of traits we obtained from supplementary datasets, excluding Sjogren's syndrome, which is absent from this table) and considering source "Omim" part of GWASdb.

| | GWASdb & OMIM | GWASdb | OMIM |
|---|---|---|---|
| Preclinical to Phase I | 1.1 (1.1-1.2) | 1.1 (1-1.1) | 1.2 (1.1-1.2) |
| Phase I to Phase II | 1.2 (1.1-1.3) | 1.2 (1.1-1.3) | 1.2 (1.1-1.3) |
| Phase II to Phase III | 1.5 (1.3-1.7) | 1.4 (1.1-1.7) | 1.6 (1.3-1.9) |
| Phase III to Approved | 1.1 (1-1.2) | 1 (0.8-1.2) | 1.1 (0.9-1.3) |
| Phase I to Phase III | 1.8 (1.5-2.1) | 1.7 (1.4-2.1) | 1.9 (1.5-2.3) |
| Phase I to Approved | 2 (1.6-2.4) | 1.7 (1.3-2.2) | 2.2 (1.6-2.8) |

Table S2: Replication of Table 1N (association between genetic evidence and historical progression) from Nelson et al. supplementary datasets. Risk ratio p(approved | genetic support)/p(approved | no genetic support) and bootstrap 95% confidence intervals.

#### S1.2 Sensitivity analysis

Many analysis decisions were made in the original publication that could affect conclusions, including the scope of genes and indications analyzed, the method of linking GWAS variants to genes, and the criteria for whether a gene target has genetic evidence for an indication. We can assess sensitivity to two key decisions using only data reported in the Nelson et al. supplementary materials.

The MeSH similarity parameter is used to dichotomize trait-indication pairs as similar or not similar. A compelling result from the original publication is that gene target-indication pairs are more likely to progress to the next stage when there is support for association of the target with a similar trait. This pattern is sensitive to the MeSH similarity cutoff, especially using GWASdb (Figure S3). The chosen cutoff 0.7 appears optimal, with more confidence limits excluding zero than at nearby values of 0.5 and 0.9. The tradeoff is expected, as lower cutoff value would be expected to include both more relevant hits and more irrelevant hits. A high proportion of irrelevant hits should lower the estimated effect size, while low numbers of total hits from a high cutoff value will lead to wide

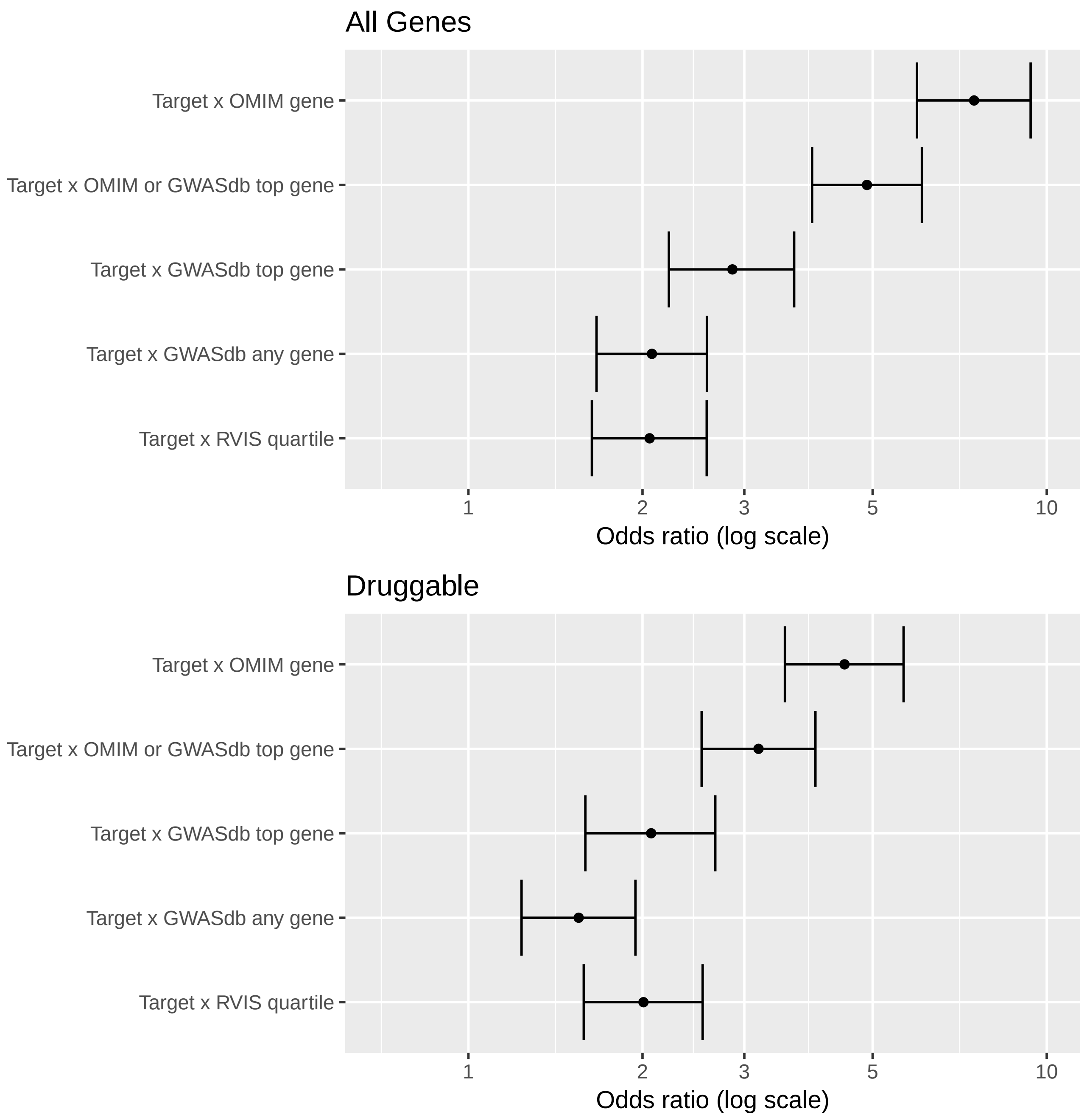

Figure S1: Replication of Figure 2N from Nelson et al. supplementary datasets. Figure shows the enrichment of approved drug targets among genes with human genetic associations.

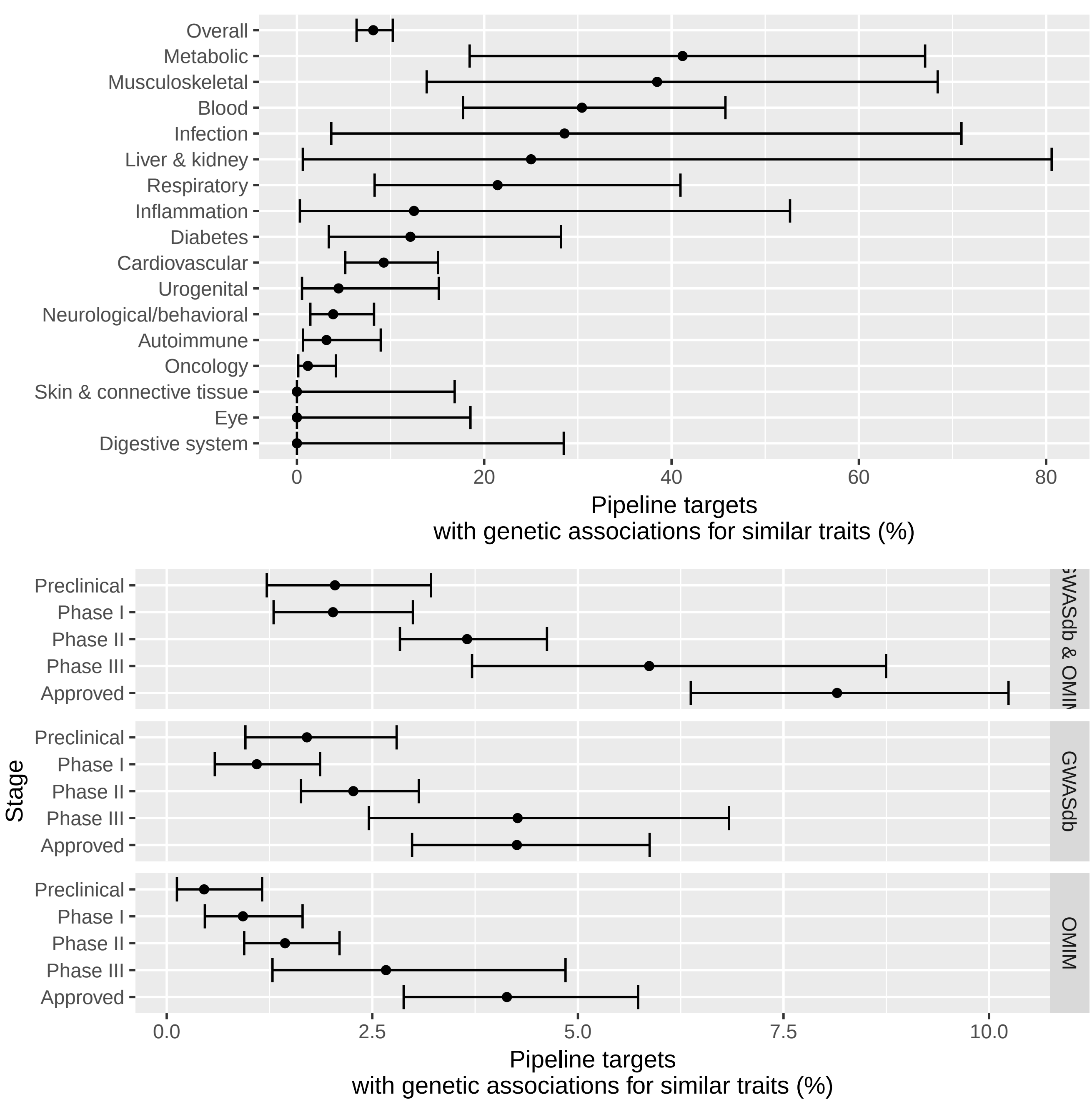

Figure S2: Replication of Figure 3N from Nelson et al. supplementary datasets. Figure shows the proportion of gene target-indication pairs with genetic associations for similar traits by pipeline phase and association source.

error bars. It is encouraging that decreasing the value to presumably include irrelevant hits removes the pattern more than increasing the value. The fact that the pattern of increasing enrichment at higher development phases for OMIM genes, but not GWAS genes, persists for unrelated traits may reflect the fact the OMIM genes are more highly enriched among approved drug targets regardless of indication (Figure S1).

Another parameter chosen in this study was the number of genetic associations required for an indication to be included in the analysis. The original study used a value of 5. The number 5 was chosen in response to the tradeoff between the number of drug indications and the percent of indications with genetic support (Supplementary Figure 7 from the original paper). We considered cutoff values between zero (no filtering on number of genetic associations) and 50 (such that only the top 19% of indications were included).

The estimated effect of genetic support on phase-specific progression probabilities is relatively insensitive to this value, though the selected value is near an optimum for the GWAS genetic evidence progression risk ratio (Figure S4). We still see a pattern of increasing enrichment of genetically supported targets in more advanced pipeline phases.

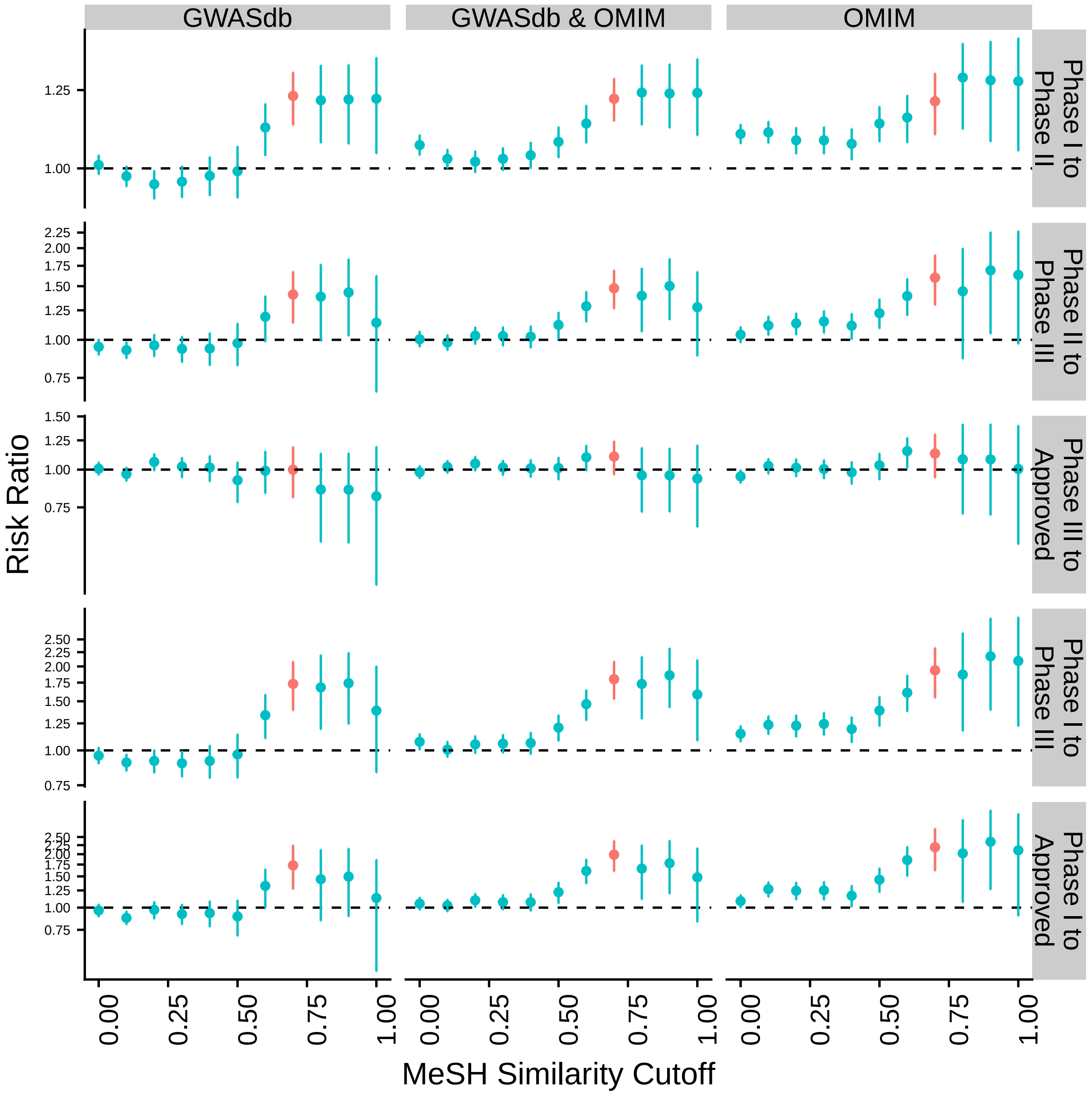

Figure S3: Sensitivity of risk ratios p(approved | genetic support)/p(approved | no genetic support) and 95% confidence limits to choice of MeSH similarity cutoff. Nelson et al. value 0.7 shown in red. Results are computed from Nelson et al supplementary datasets.

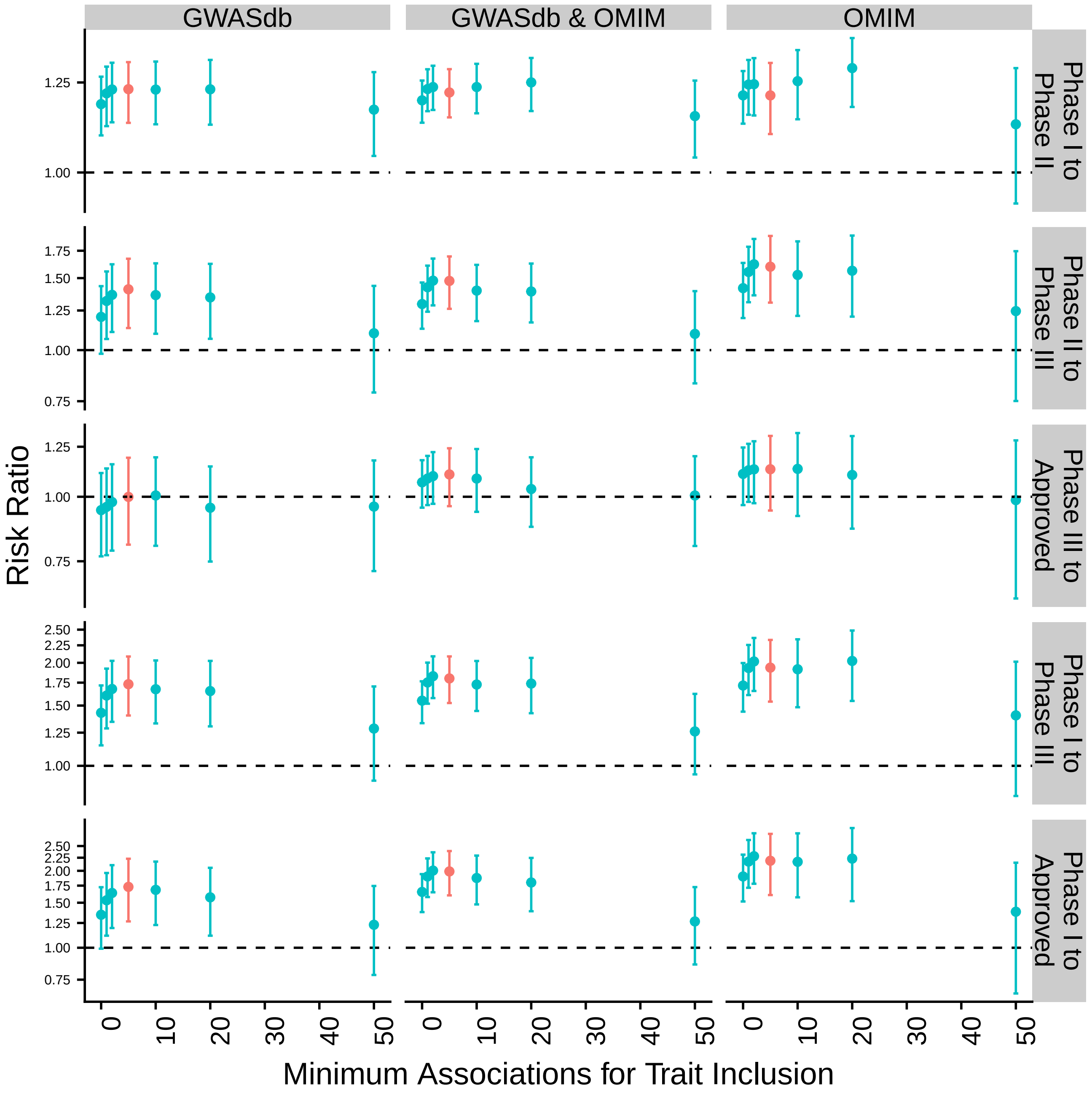

Figure S4: Sensitivity of risk ratios p(approved | genetic support)/p(approved | no genetic support) and 95% confidence limits to choice of minimum number of associations parameter. Nelson et al. value of 5 shown in red. Results are computed from Nelson et al supplementary datasets.

### S2 Updating Pipeline Data: Supplementary Methods And Results

#### S2.1 Updating Pipeline Data with New Pharmaprojects Data Freeze

Informa Pharmaprojects [1] data were obtained from an XML file of the full database from Jan 25, 2018. Following Nelson et al., we excluded drugs with nonhuman or xMHC gene targets. We considered xMHC to include HIST1H2AA and KIFC1 and all genes between them (Chromosome 6 25.7 Mb-33.4 Mb). We also excluded drugs with non protein-coding gene targets. All Entrez ids were mapped to ensembl ids and indications to MeSH headings. Unmapped indications and gene targets were excluded. The drug table was collapsed to one row per gene target-MeSH pair. The latest phase of gene target-MeSH pair $(g, m)$ is defined to be the latest phase of any indication mapping to MeSH heading $m$ over all drugs with target $g$. $(g, m)$ is US-EU approved if there exists a drug $d$ satisfying $d$ is US/EU approved, $d$ is approved for an indication mapping to MeSH heading $m$, and $d$ has target $g$.

Processing Pharmaprojects data as outlined above does not deviate greatly from the approach used by Nelson et al. However, we made one major methodological change in how latest phase is assigned to Pharmaprojects drug-indication pairs. 66% of drug-indication pairs with human gene targets in the Pharmaprojects database are assigned an inactive status such as No Development Reported, but determining the effect of genetic evidence on clinical development progression requires the latest development phase attained by each gene target-indication pair. For this reason, inactive gene target-indication pairs were excluded from key analyses in [20]. We were concerned that excluding these drugs would bias the analysis as active drug-indication pairs have been under development for shorter time periods, on average, and therefore may not have had sufficient time to become approved. We assigned a latest historical development phase to these pairs using other fields of the Pharmaprojects database whenever possible. Our assignment procedure is detailed in the next section.

##### S2.1.1 Pharmaprojects latest phase assignment: Methods

We attempt to assign each drug and each drug-indication pair a latest historical phase that is one of "Preclinical", "Phase I Clinical Trial", "Phase II Clinical Trial", "Phase III Clinical Trial", "Pre-registration" or "Approved" (in order of earliest to latest development phase). "Approved" includes drugs with status "Launched", "Registered", or "Withdrawn" in Pharmaprojects. The latest historical development phase is the most advance phase of development a drug-indication pair has reached in any point of its history. Drugs or drug-indication pairs with status "Discontinued", "No Development Reported", "Not Applicable", "Suspended", NA, or "-" are considered to have *unknown latest phase*, and we would like to assign them a known one. We do this using other fields in the Pharmaprojects database. To distinguish between our inferred latest phase and raw Pharmaprojects data fields, we will refer to the former as the *latest phase*, and the latter as a status (disease status, global status, or country status, depending on which field is used). We will also distinguish the *global latest phase* (or status), the latest phase of a drug for any indication, and the latest phase at the indication level. Because all the main analysis of the paper are performed using indication-level phases, a phase is measured at the indication level unless otherwise specified.

Data from the event history, country status, clinical information, global status, and disease status for each drug was used in determining latest phase. The general strategy will be to find a latest phase using several different sources of information, then assign the most advanced phase over all the sources.

###### Assigning Phase From Event History

Many Pharmaprojects entries have an event history including changes in global and disease status. These fields have a standardized set of event types, and fairly consistent formatting of event details. Some events can be assigned a phase. Any event with event type "First Launches" or "Additional Launches" is assigned phase launched, events of type "Registration Submissions" are assigned the "Pre-registration" phase, events of type "First Registrations" or "Additional Registrations" are assigned the "Registration" phase, and event type "Withdrawn Products" is assigned the withdrawn phase. Trial phases in event histories were standardized to eliminate letters (e.g Phase Ia to Phase I) and to change combined phase trials to the later of the two phases (e.g. Phase II/III to Phase III). Clinical trial phases were found in event details using string pattern matching. Global highest phase for the event history source was found by taking the latest phase of any event for that drug. Latest indication-level phase from event history was taken to be the latest phase for any event involving that drug in which the disease name also appeared in the event description.

##### Assigning Phase From Country Status

Nearly all drugs have a known development status in some country (Table S3). Unlike global status and disease status, country status almost never reverts to an inactive status such as "No Development Reported", "Discontinued" or "Suspended" (inferred from the rarity or absence of these terms). Latest global phase was determined from the country status field as the top phase for a drug over any country. Unfortunately, country status cannot be used to determine indication-level phases as it does not contain any indication information.

##### Assigning Phase From Clinical Trial Information

Pharmaprojects contains text descriptions of Preclinical, Phase I, Phase II, and Phase III trials in drug entries. These entries are unstructured and are (usually) empty when no trials of that phase have been performed. Therefore the presence of text in these fields is closely related to whether trials have been performed for that phase, though these fields may include trials that were planned but never completed. To reduce the possibility of error, we only used these fields to assign a phase when the text contained the name of the phase and did not contain the words "planned" or "expected." The latest phase with respect to the trial descriptions was the top trial phase with description field satisfying the required conditions. To assign preclinical phase, we only required this field have more than 5 characters (to eliminate a small number of nonsense entries). Sometimes these fields contain one or more Pharmaprojects indication terms formatted to match Pharmaprojects drug indications. The latest indication phase for a drug was assigned to be the latest stage field containing that indication term, subject to the above mentioned restrictions for quality control.

##### Determining Global Latest Phase From Other Fields

We have computed the latest phase of Pharmaprojects drugs using several fields. We can use this to create the global latest phase field, giving the latest known historical pipeline phase. Data from the country status and event fields are only rarely discordant with known global status (in the sense that they infrequently imply a latest phase different from the Pharmaprojects global status when the latter is a clinical phase, Table S4). Sometimes, discordance reflects status reversions in which development ceases and then restarts at lower phase, and the event history may actually more reliably give the most advanced development phase attained by the drug. Data from the clinical trial phase description fields tend to be most discordant. Given these different degrees of error, the global latest phase for a drug was determined as follows

1. Latest phase is taken to be the most advanced development phase using the Pharmaprojects global status, country status and event status. 98.9% of drugs had a latest global phase assigned in this manner.
2. Those global latest phases that are still unknown are determined from the population of the clinical trial text fields.
3. Remaining unassigned global latest phases are retained at their original value. In all, 99.4% of drugs could be assigned a latest phase.

##### Using Global Latest Phase to Determine Indication Latest Phase

We used the global latest phase of a drug to assign a indication latest phase with some simple assumptions.

1. When the global latest phase is Preclinical, assume all indications are in the Preclinical phase (Pharmaprojects does not have a category lower than Preclinical).
2. When there is only one indication, the latest phase for the indication is the same as the global latest phase (assumes no indications have been omitted).

With these definitions, we assigned global latest phase whenever possible, creating a more complete dataset for evaluation (Figure S5).

##### S2.1.2 Pharmaprojects Latest Phase Assignment: Results

##### Assigned Status By Date

99.4% of drugs and 84.6% of drug-indication pairs can be assigned a latest phase. This reduces the dramatic differences in last modified date (Figure S6) and in the date of the first recorded drug event (Figure S7) between drugs in clinical trials and approved drugs as compared to the approach of excluding these results used in [20]. Although reduced, there are still systematic differences in dates between drugs with unknown latest phase and other drugs.

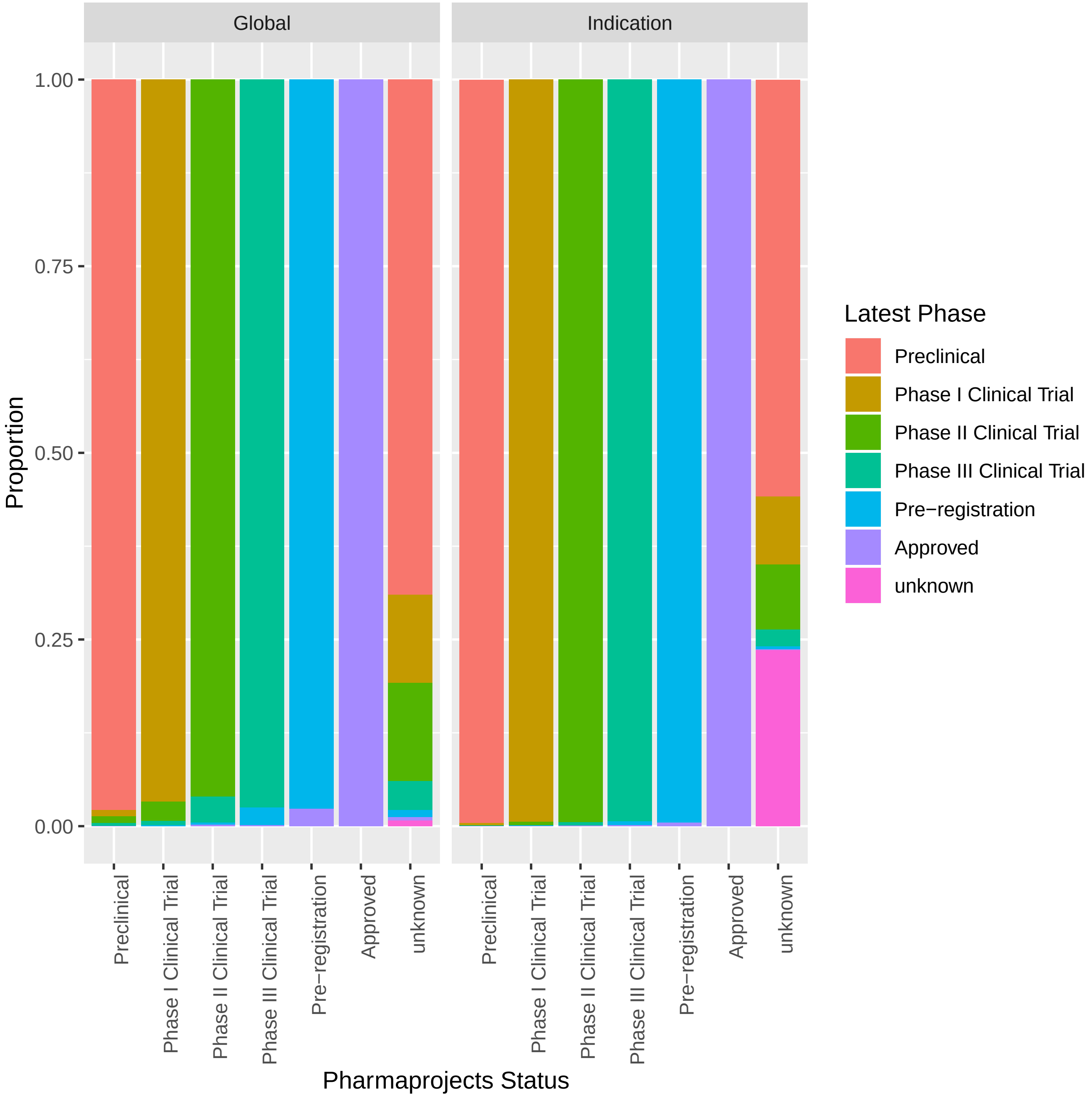

Figure S5: Assigned latest phase compared to Pharmaprojects status (unknown Pharmaprojects status categories such as No Development Reported and Suspended are combined) at the global and indication level.

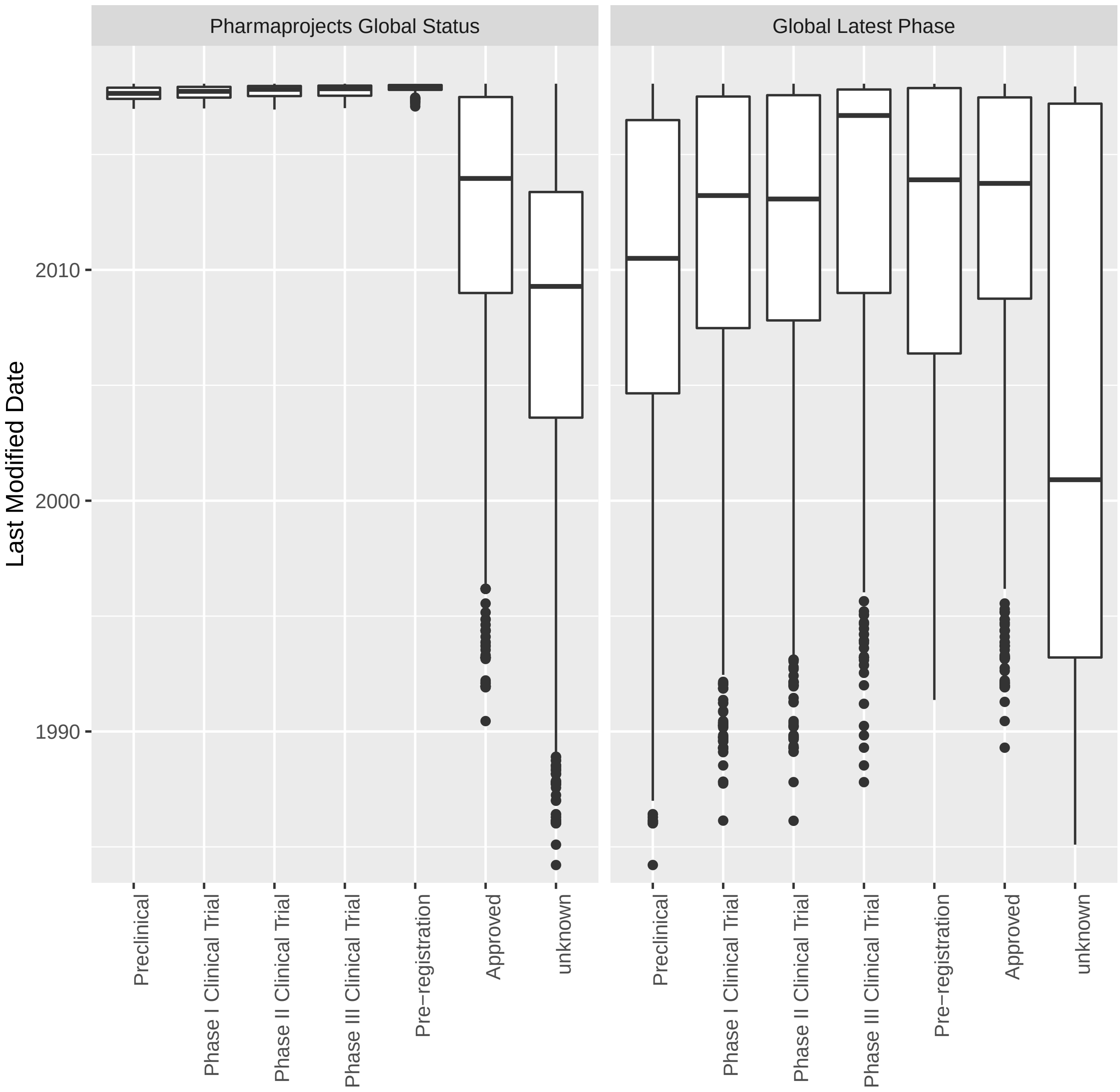

Figure S6: Latest change date to Pharmaproject entry for drugs by development status. In panel Pharmaprojects Global Status, statuses come from the Pharmaprojects global status field. In panel Global Latest Phase, statuses are the latest global development phase assigned in this document.

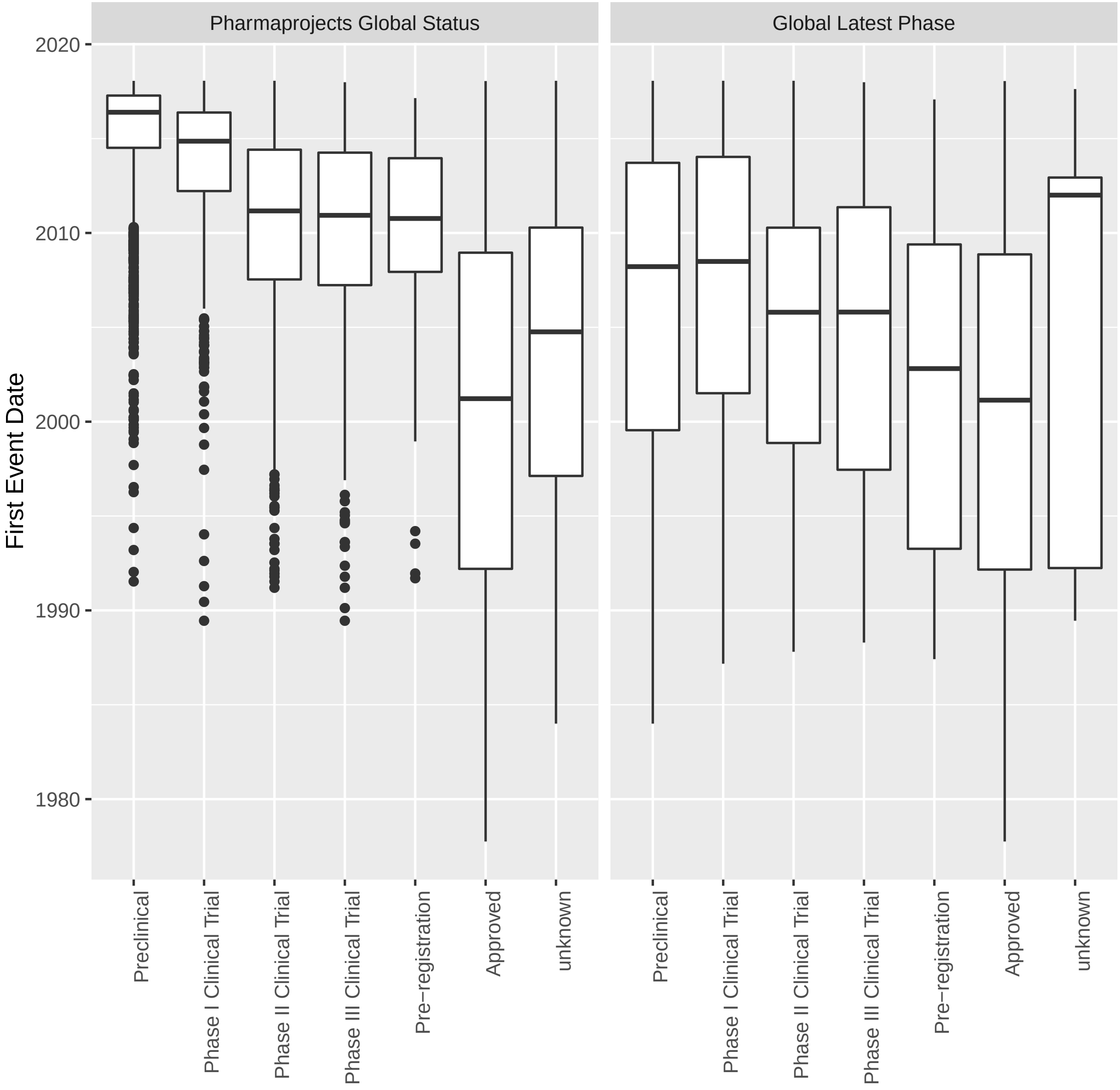

Figure S7: First event date for Pharmaproject entry for drugs by development status. In panel Pharmaprojects Global Status, statuses come from the Pharmaprojects global status field. In panel Global Latest Phase, statuses are the latest global development phase assigned in this document. Note 6% of compounds do not have any entries in their event history and are omitted.

**Availability and Quality of Data by Source**

When disease or global status is a known phase, it usually matches the latest phase assigned by our procedure (Figure S5). Drug-indication pairs with inactive disease statuses (with unknown pipeline phase) tend to have lower latest phase than drugs in the dataset as a whole, with preclinical pairs more common and approved drug-indication pairs rare.

Pharmaprojects fields differ in the proportion of drugs or drug-indication pairs for which they are informative. Country status is the most readily available, while global status (for single indication and preclinical compounds) is the most informative source of data on indication-level phase (Table S3). Both of these sources of information are highly internally consistent with Pharmaprojects global and disease status when both are a known trial phase (Table S4). The event history is also usually internally consistent with Pharmaprojects assigned status, and commonly implies a global, but rarely an indication-level phase. Due to the sparsity of indication status information available in the event history, it is not surprising that where this information differs from the Pharmaproject disease status, it tends to be a lower status. Assigning status from the clinical trial information field shows the least agreement with global status, and there is a strong tendency for assignments using this method to have a lower trial phase, perhaps due to the filtering procedure inappropriately excluding records or due to missing information. Assignments made solely on the basis of parsing this field will be biased towards being lower than the latest development status (Table S4).

| Source | Indication | Global |
|---|---|---|
| Event | 0.06 | 0.58 |
| Global | 0.54 | |
| Info | 0.17 | 0.74 |
| Country | | 0.98 |

Table S3: Proportion of drug-indication pairs (Indication) or drugs (Global) having development status information available from each source. Event=Pharmaprojects event history, Info=Pharmaprojects clinical information fields, Country=Pharmaprojects country status, Global=Indication status inferred from global status.

| Source | Type | Less | Greater | Equal |
|---|---|---|---|---|
| Global | Indication | 0.00 | 0.00 | 1.00 |
| Country | Global | 0.01 | 0.00 | 0.99 |
| Event | Global | 0.01 | 0.02 | 0.97 |
| Event | Indication | 0.08 | 0.02 | 0.91 |
| Info | Global | 0.46 | 0.02 | 0.53 |
| Info | Indication | 0.50 | 0.02 | 0.48 |

Table S4: Agreement between Pharmaproject status (Type=Global or Indication) and latest phase using each evidence source when both are assigned a known development status. Columns less, greater, and equal are the proportion of times in which the source implicates a latest pipeline phase less advanced than, more advanced than, or equal to that reported by Pharmaprojects. Arranged in order of descreasing agreement.

#### S2.2 Supplementary Results Using Updated Pipeline Data

##### S2.2.1 Analysis of All Updated Pipeline Data

| | GWASdb & OMIM | GWASdb | OMIM |
|---|---|---|---|
| Preclinical to Phase I | 1.1 (1.1-1.2) | 1.1 (1-1.1) | 1.2 (1.1-1.4) |
| Phase I to Phase II | 1.1 (1-1.2) | 1.1 (1-1.1) | 1.2 (1.1-1.2) |
| Phase II to Phase III | 1.4 (1.3-1.6) | 1.2 (1-1.4) | 1.8 (1.5-2) |
| Phase III to Approved | 1.2 (1.1-1.3) | 1.2 (1-1.3) | 1.3 (1.1-1.4) |
| Phase I to Phase III | 1.6 (1.4-1.8) | 1.3 (1-1.5) | 2.1 (1.8-2.4) |
| Phase I to Approved | 2 (1.6-2.3) | 1.5 (1.1-1.9) | 2.6 (2.1-3.2) |

Table S5: Replication of Table 1N (association between genetic evidence and historical progression) from Nelson et al. supplementary genetic association dataset and updated pipeline data. Risk ratio p(approved | genetic support)/p(approved | no genetic support) and bootstrap 95% confidence intervals.

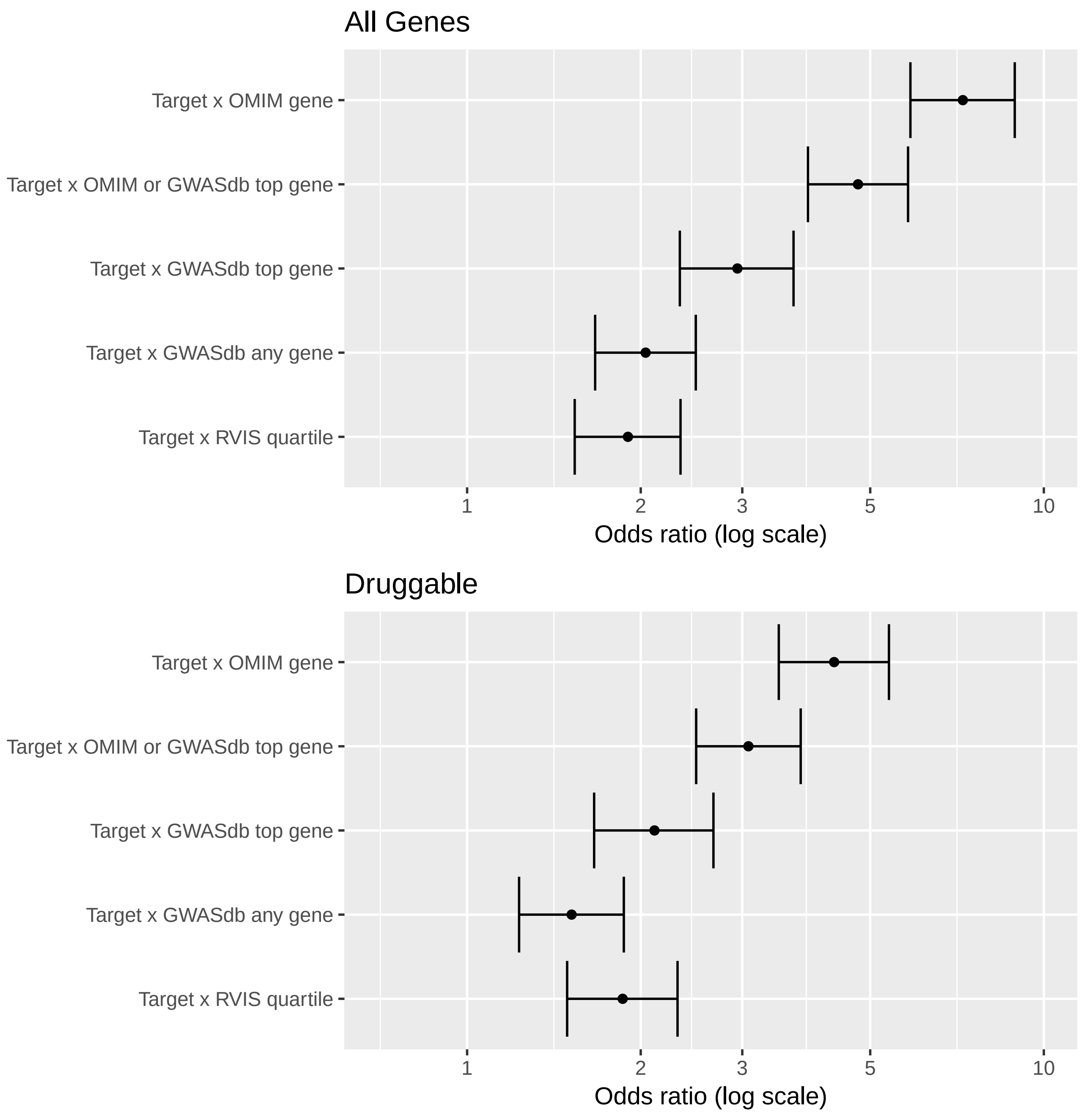

Figure S8: Replication of Figure 2N from Nelson et al. supplementary genetic association dataset and updated pipeline data. Figure shows the enrichment of approved drug targets among genes with human genetic associations.

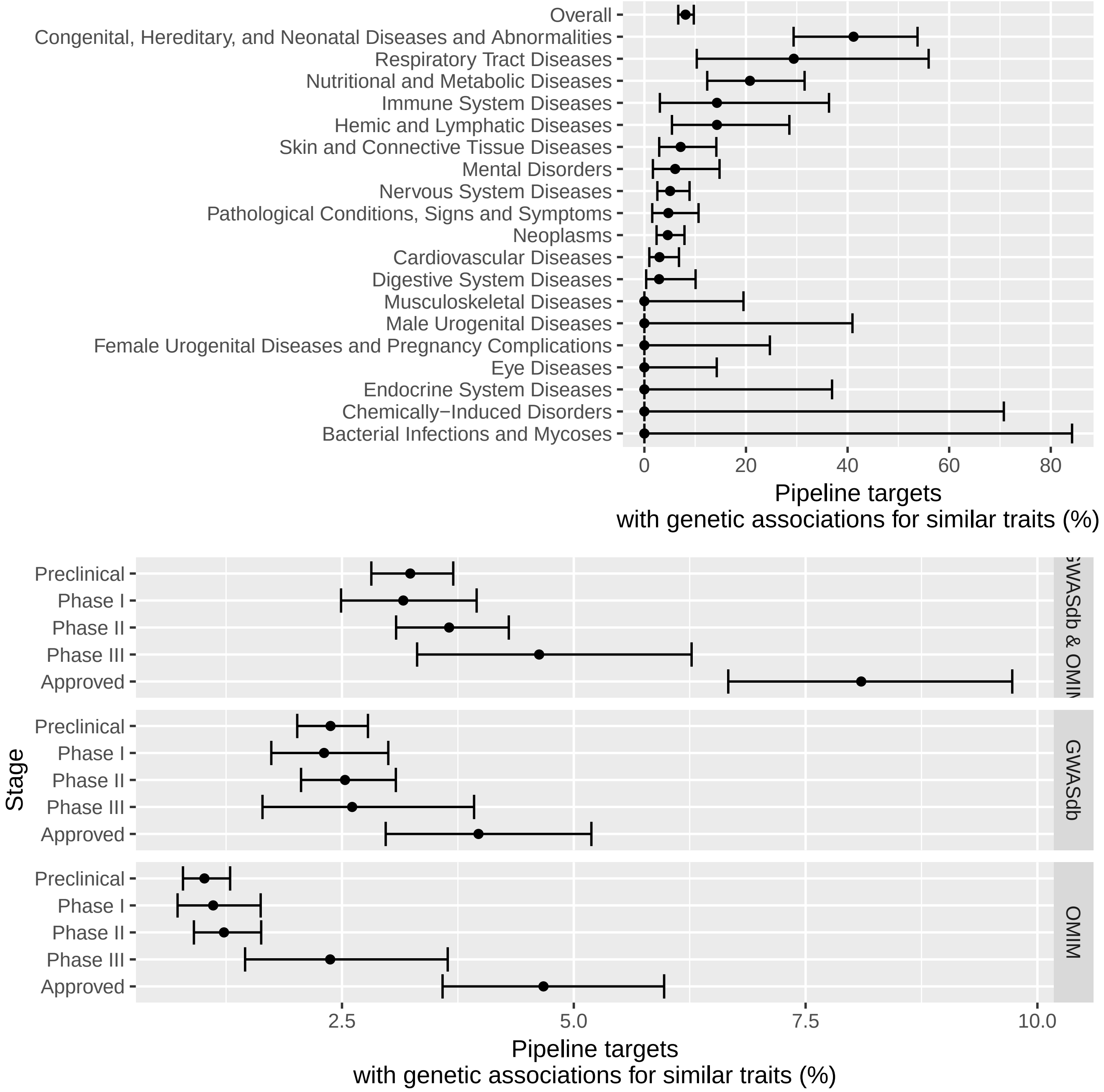

Figure S9: Replication of Figure 3N from Nelson et al. supplementary genetic association dataset and updated pipeline data. Figure shows the proportion of gene target-indication pairs with genetic associations for similar traits by pipeline phase and association source.

##### S2.2.2 Analysis of 2013-2018 Progressions (Pipeline Progression)

| Event | GWASdb & OMIM | GWASdb | OMIM | N |
|---|---|---|---|---|
| Preclinical to Phase I | 1.7 (0.8-2.6) | 1.8 (0.8-2.9) | 1.8 (0.5-3.2) | 829 (207) |
| Phase I to Phase II | 1.6 (1.1-2.2) | 1.5 (0.7-2.2) | 1.9 (1.1-2.7) | 986 (362) |
| Phase II to Phase III | 1.6 (0.9-2.3) | 0.4 (0-1) | 2.8 (1.7-4.1) | 1532 (250) |
| Phase III to Approved | 1.5 (0.8-2.3) | 1.5 (0.6-2.4) | 1.2 (0.3-2.3) | 341 (125) |

Table S6: Risk ratio of pipeline progression from 2013 to 2018 by presence or absence of supporting genetic evidence and 2013 phase. Risk ratio and 95% confidence intervals. Last column gives the total number of gene target-indication pairs labeled with that phase in 2013 and the total number of gene target-indication pairs that progressed in development.

| Event | GWASdb & OMIM | GWASdb | OMIM | N |
|---|---|---|---|---|
| Progression | 1.5 (1.1-1.8) | 1.2 (0.8-1.6) | 1.9 (1.4-2.4) | 3842 (961) |
| Approval | 2.5 (1.4-4) | 2.5 (1.1-4.1) | 2.5 (1-4.6) | 3842 (172) |

Table S7: Risk ratio of pipeline progression from 2013 to 2018 by presence or absence of supporting genetic evidence. Risk ratio and 95% confidence intervals. Last column gives the total number of 2013 gene target-indication pairs included in the analysis and the total number of drugs that progressed in development. This table shows the risk ratio of progression to a higher phase or to approval from any starting phase.

###### S2.2.3 Analysis of Previously Unused Gene Target-Indication Pairs (New Pipeline)

| | GWASdb & OMIM | GWASdb | OMIM |
|---|---|---|---|
| Preclinical to Phase I | 1.1 (1-1.2) | 1.1 (1-1.2) | 1.1 (0.9-1.3) |
| Phase I to Phase II | 1 (0.9-1.1) | 1 (0.9-1.2) | 0.9 (0.7-1.1) |
| Phase II to Phase III | 1.3 (0.8-1.7) | 1.3 (0.8-1.8) | 1.3 (0.6-2.2) |
| Phase III to Approved | 1.9 (1.1-2.7) | 1.7 (0.9-2.5) | 2.8 (1.5-3.9) |
| Phase I to Phase III | 1.3 (0.8-1.8) | 1.3 (0.8-1.9) | 1.2 (0.5-2.1) |
| Phase I to Approved | 2.4 (1.2-3.9) | 2.2 (0.9-3.8) | 3.4 (1.1-6.6) |

Table S8: Replication of Table 1N (association between genetic evidence and historical progression) from Nelson et al. 2015 supplementary genetic association dataset and updated pipeline data, using only gene target-indication pairs not used in the original analysis either due to not being in the table of gene target-indication pairs or having an inactive development status. Risk ratio p(approved | genetic support)/p(approved | no genetic support) and bootstrap 95% confidence intervals.

| | GWASdb & OMIM | GWASdb | OMIM |
|---|---|---|---|
| Preclinical to Phase I | 1.2 (1-1.4) | 1.3 (1.1-1.5) | 0.9 (0.6-1.2) |
| Phase I to Phase II | 1 (0.9-1.2) | 1 (0.9-1.2) | 1 (0.7-1.4) |
| Phase II to Phase III | 1.6 (1-2.4) | 1.4 (0.8-2.3) | 3 (1.4-4.7) |
| Phase III to Approved | 1.5 (0.7-2.5) | 1.4 (0.5-2.5) | 2.3 (1-3.7) |
| Phase I to Phase III | 1.7 (1-2.5) | 1.5 (0.7-2.4) | 3 (1-5.2) |
| Phase I to Approved | 2.5 (0.9-4.7) | 2.1 (0.4-4.3) | 7 (1.6-14.3) |

Table S9: Replication of Table 1N (association between genetic evidence and historical progression) from Nelson et al. 2015 supplementary genetic association dataset and updated pipeline data, using only gene target-indication pairs not used in the original analysis due to not being in the table of gene target-indication pairs. Risk ratio p(approved | genetic support)/p(approved | no genetic support) and bootstrap 95% confidence intervals.

| | GWASdb & OMIM | GWASdb | OMIM |
|---|---|---|---|
| Preclinical to Phase I | 1.1 (0.9-1.3) | 0.9 (0.7-1.1) | 1.4 (1.1-1.8) |
| Phase I to Phase II | 1 (0.8-1.2) | 1 (0.8-1.2) | 0.9 (0.6-1.2) |
| Phase II to Phase III | 0.9 (0.5-1.5) | 1.2 (0.5-2) | 0.5 (0-1.2) |
| Phase III to Approved | 2.5 (1.1-3.9) | 2 (0.5-3.5) | 4 (3.3-5) |
| Phase I to Phase III | 0.9 (0.4-1.5) | 1.2 (0.5-2) | 0.4 (0-1) |
| Phase I to Approved | 2.3 (0.7-4.4) | 2.4 (0.5-5.1) | 1.6 (0-4.3) |

Table S10: Replication of Table 1N (association between genetic evidence and historical progression) from Nelson et al. 2015 supplementary genetic association dataset and updated pipeline data, using only gene target-indication pairs not used in the original analysis due to being assigned an inactive status. Risk ratio p(approved | genetic support)/p(approved | no genetic support) and bootstrap 95% confidence intervals.

###### S2.2.4 Removing similar mechanisms to 2013 Approved Drugs

Target-indication pairs supported by genetic evidence were more likely to progress from 2013-2018, and the effect was statistically significant for OMIM drugs. Nelson et al. could not have been aware of progressions that occured after the creation of their dataset, so we reason these results should be minimally affected by any overfitting to the original dataset. However, progressions from 2013-2018 may not actually be independent of pre-2013 approvals, because approved drugs may be repurposed for other indications. Target-indication pairs with similar but not identical indications might be expected to have both positively correlated approvals and positively correlated genetic evidence.

We address this concern by labelling each target-indication pair with its similarity to US/EU approved target-indication pairs in the original Nelson et al. dataset downloaded in 2013. Specifically, let $\mathcal{U}$ be the set of US/EU approved target-indication pairs in the original dataset. Define $S_A((t,g)) = \max_{p\in\mathcal{U}} S((t,g),p)$ where $S$ is the target-indication similarity function defined in Section 5.4 of the main text. We will say $(t,g)$ has a similar 2013 approved pair if $S_A(t, g) \geq 0.73$ (our recalibrated version of the Nelson et al. cutoff of 0.7). We will eliminate all such pairs from the progression dataset and recompute the effect of genetic evidence on phase progression.

As predicted, we find that 2013-2018 progression is positively associated with $S_A$, the trait similarity of the most similar 2013 approved target-indication pair sharing the target (not shown).

| Event | GWASdb & OMIM | GWASdb | OMIM | N |
|---|---|---|---|---|
| Preclinical to Phase I | 1.9 (1-2.8) | 1.8 (0.8-2.9) | 2.3 (1-3.8) | 818 (202) |
| Phase I to Phase II | 1.7 (1.1-2.3) | 1.5 (0.8-2.3) | 2 (1.1-2.7) | 968 (350) |
| Phase II to Phase III | 1.6 (0.9-2.4) | 0.4 (0-1) | 3.2 (1.9-4.7) | 1442 (231) |
| Phase III to Approved | 1.6 (0.9-2.4) | 1.5 (0.7-2.5) | 1.3 (0.3-2.5) | 289 (102) |

Table S11: Risk ratio of progression in clinical development from 2013 to 2018 by presence or absence of supporting genetic evidence. Calculations are performed on the subset of target-indication pairs without similar 2013 approved target-indication pairs, using similarity cutoff 0.73. Risk ratio and 95% confidence intervals.

However, estimated effects of genetic evidence that were previously positive remain positive, and the positive effect of OMIM genetic evidence on Phase II to Phase III transitions increases and remains significant when target-indication pairs with similar 2013 approved target-indication pairs are removed from the dataset (Table S11). In fact, these patterns are largely maintained even when excluding all targets with an approved indication in 2013 (Table S12).

| Event | GWASdb & OMIM | GWASdb | OMIM | N |
|---|---|---|---|---|
| Preclinical to Phase I | 2.1 (1-3.2) | 2 (0.7-3.3) | 2.3 (0.7-4.2) | 633 (140) |
| Phase I to Phase II | 1.6 (0.9-2.5) | 1.7 (0.7-2.7) | 1.8 (0.5-2.9) | 655 (215) |
| Phase II to Phase III | 2.2 (1-3.5) | 0.3 (0-1.1) | 5.5 (3.4-7.8) | 754 (106) |
| Phase III to Approved | 0.9 (0.3-1.6) | 1.1 (0.4-2) | 0.4 (0-1.2) | 100 (45) |

Table S12: Risk ratio of progression in clinical development from 2013 to 2018 by presence or absence of supporting genetic evidence. Calculations are performed on the subset of target-indication pairs with no approved 2013 drugs for that target. Risk ratio and 95% confidence intervals.

##### S2.2.5 Additional Replication Set: OMIM supplementary concepts

Our reanalysis of OMIM allows for one additional replication set. Mendelian disorders often appear in the MeSH vocabulary as supplementary concepts. 79% of OMIM traits, but no Pharmaprojects indications, were mapped to MeSH supplementary concepts. Supplementary concepts were assigned zero similarity to all drug indications because they are not part of the hierarchical structure of the MeSH vocabulary used to compute similarities (see S4.1), and therefore did not constitute supporting genetic evidence for any drug mechanism. OMIM disorders mapping to MeSH supplementary concepts can be assigned to mapped headings supplied by MeSH. When there were multiple mapped headings, disease and psychiatric terms were preferred, and terms with higher information content were also preferred (measured by fewer descendants in the MeSH hierarchy, see S4.1).

Estimated risk ratios of historical progression are in Table S13. The positive effect of OMIM genetic evidence remains when mapped headings are used to identify indications similar to OMIM traits mapped to MeSH supplementary concepts.

| | All OMIM | New OMIM Only |
|---|---|---|
| Preclinical to Phase I | 1.1 (1.1-1.2) | 1.1 (0.9-1.2) |
| Phase I to Phase II | 1.2 (1.1-1.3) | 1.2 (1.1-1.3) |
| Phase II to Phase III | 1.6 (1.4-1.8) | 1.4 (1.1-1.8) |
| Phase III to Approved | 1.3 (1.2-1.4) | 1.3 (1.1-1.5) |
| Phase I to Phase III | 1.9 (1.7-2.2) | 1.7 (1.3-2.2) |
| Phase I to Approved | 2.5 (2.1-2.9) | 2.2 (1.6-2.9) |

Table S13: Replication of Table 1N (association between genetic evidence and historical progression) from Nelson et al. supplementary genetic association dataset and updated pipeline data with all MeSH terms assigned valid headings (All OMIM). Column New OMIM shows results using only OMIM entries originally mapped to supplementary concepts. Risk ratio p(approved | genetic support)/p(approved | no genetic support) and bootstrap 95% confidence intervals.

| Dataset or Tool | Downloaded | Version | Filter | URL or file path |
|---|---|---|---|---|
| GWAS Catalog | Sept 25, 2017 | v1.0.1 | $p < 10^{-8}$ | https://www.ebi.ac.uk/gwas/api/search/downloads/alternative |
| 1000 Genomes | | Phase 3 | $r^2 \geq 0.5$ | bochet.gcc.biostat.washington.edu/beagle/1000_Genomes_phase3_v5a |
| GTEx | Sept 26, 2017 | v7 | $p < 10^{-6}$ | https://www.gtexportal.org/home/datasets, GTEx_Analysis_v7_eQTL.tar.gz |
| SnpEff | | | $d \leq 5000$ | http://snpeff.sourceforge.net/ |
| Regulatory Elements Database | Oct 4, 2017 | v3 | $p > 0.999$ | http://big.databio.org/RED/allGeneCorrelations100000.p2_v3.txt.gz |
| RegulomeDB | Oct 23, 2017 | v1.1 | | http://www.regulomedb.org/downloads/RegulomeDB.dbSNP141.txt.gz |

Table S14: Sources used to obtain and annotate GWAS variants. Filter gives the $p$-value or other cutoff used to filter results.

### S3 Updated Genetic Dataset: Supplementary Methods and Results

#### S3.1 Updating GWAS Dataset with GWAS Catalog and GTEx

##### S3.1.1 Methods

**Nelson et al. Gene Table Creation and Sources** Nelson et al. obtained SNP-trait associations primarily from GWASdb [14]. The list of SNPs was expanded to any SNP with $r^2 \geq 0.5$ with the reported SNP. These SNPs will be referred to as LD SNPs. LD SNPs were mapped to genes on the basis of three criteria. LD SNPs within 5 kb upstream or downstream of the gene were mapped to that gene. LD SNPs with eQTLs for the gene were mapped to that gene. Finally, LD SNPs with a DNase Hypersensitive Site (DHS) correlated with the transcription start site of the gene were mapped to that gene [18].

**Recreating Gene Table with Updated Sources**

**Obtaining Datasets** Table S14 gives URLs and download dates for all major data sources and filtering criteria used if applicable. Cutoff values were chosen to match those in Nelson et al. when available. Genetic association data was taken from the GWAS Catalog [16]. The LD SNP expansion was conducted in Plink [23] using the 1000 Genomes Project [5] Phase 3 in the EUR population. eQTL associations were GTEx significant eQTL in any tissue ($p < 10^{-6}$). DHS-gene correlations were taken from the Regulatory Element Database [26], which reported DHS activity correlations with gene expression and one-sided $p$-values. The RegulomeDB [3] was downloaded Oct 23, 2017. SnpEff [4] was used find genes within 5 kb of each LD SNP in hg38 and annotate potentially deleterious variant LD SNPs, including missense variants.

**Linking Datasets** GWAS Catalog, 1000 Genomes, and the RegulomeDB supplied SNP rs ids. GTEx variant rs ids were obtained from GTEx (GTEx_Analysis_2016-01-15_v7_WholeGenomeSeq_635Ind_PASS_AB02_GQ20_HETX_MISS15_PLINKQC.lookup_table.txt, downloaded from `https://gtexportal.org/home/datasets`). SnpEff required a vcf file as input. A vcf file for all human short variations from dbSNP build 150 was obtained from `ftp://ftp.ncbi.nih.gov/snp/organisms/human_9606/VCF/00-All.vcf.gz` and filtered to include only SNPs of interest using `vcftools` [7]. The filtered file was input to SnpEff. Gene symbols from SnpEff were mapped to ensembl ids.

**Scoring Genes** The above annotation procedure created datasets with pairs of LD SNPs and genes linked by eQTL, distance, or DHS correlation evidence. Additionally, each LD SNP can be annotated with its RegulomeDB score. This LD SNP annotation information was joined with a data frame of all significant SNP-trait pairs from the GWAS Catalog and a data frame linking LD SNPs and original GWAS SNPs. Gene scores (needed only for replication of Figure 2N [20], not used in main text analysis) were computed as described in [20]. When multiple LD SNPs for a GWAS hit implicated the same gene, only the highest scoring hit was retained. Finally, the dataset was filtered to retain only those genes annotated with protein coding ensembl biotype.

###### Mapping EFO Terms

GWAS Catalog traits are given both as a text description (disease trait) and as mapped EFO terms [17]. Linking with other sources requires a common ontology, in this case the NLM MeSH vocabulary. Using a combination of manual and automated procedures, we mapped each GWAS Catalog EFO term to MeSH. Many GWAS Catalog EFO terms describe measurements. Where possible, we mapped terms to a MeSH term describing a measurement. When not possible, we mapped to the entity being measured. For example, "bone density" maps to Bone Density but "eosiniphil count" maps to Eosiniphils.

An additional complication is that many reported genetic associations are mapped to multiple traits in EFO. We automatically assigned entries with multiple EFO terms using these rules where possible, and manually mapped additional terms.

1. Entries with a single distinct mapped MeSH heading are mapped to that heading.
2. If there are two mapped MeSH headings and one is ancestor of the other, map to the descendant.
3. Exclude EFO terms containing "blood metabolite measurement", "clinical laboratory measurement", "cerebrospinal fluid biomarker measurement", "Alzheimer's disease biomarker measurement", "cardiovascular measurement", "liver enzyme measurement" and assign remaining mapped MeSH term (usually a more specific measurement or measured entity).
4. Exclude EFO terms containing "onset", "survival" and assign remaining mapped MeSH term (usually a disease)
5. If the disease trait is $x$ in $y$ (e.g. Body mass index in physically active individuals) or $x$ adjusted for $y$ (e.g. Visceral adipose tissue adjusted for BMI), map to $x$.
6. If only one mapped MeSH heading is a disease, map to the disease.
7. Exclude entries with disease traits containing "combine", "interaction", " and ", "pleiotropy", "multi-trait" or " or " unless otherwise mapped.
8. Exclude entries with mapped EFO term "response to" (these are treatment response association studies).

17% of reported associations were excluded due to these rules or to lacking a satisfactory MeSH heading.

##### S3.1.2 Results

| Property | Previous | Current | Overlap |
|---|---|---|---|
| Ensembl Genes | 6045 | 8872 | 4255 |
| MeSH | 434 | 604 | 232 |
| Associated SNPs | 13933 | 10815 | 2732 |
| LD SNPs | 16270 | 18204 | 1562 |
| LD SNP-Gene Links | 20313 | 27208 | 1304 |
| SNP-Gene Links | 53526 | 48564 | 6678 |
| MeSH-Gene Links | 13702 | 36044 | 3535 |
| MeSH-Gene-SNP Links | 65923 | 62043 | 4325 |

Table S15: Comparison of counts of distinct genes, traits (MeSH), and SNPs reported by Nelson et al. and those from the current analysis. Overlap is the number of items in common.

| Property | Previous | Current | Overlap |
|---|---|---|---|
| Ensembl Genes | 3424 | 4020 | 2338 |
| MeSH | 334 | 345 | 199 |
| Associated SNPs | 3353 | 2690 | 1722 |
| LD SNPs | 5310 | 5742 | 787 |
| LD SNP-Gene Links | 6533 | 8467 | 674 |
| SNP-Gene Links | 10637 | 11741 | 4130 |
| MeSH-Gene Links | 6574 | 10316 | 2476 |
| MeSH-Gene-SNP Links | 11769 | 14139 | 2983 |

Table S16: Comparison of counts of distinct genes, traits (MeSH), and SNPs reported by Nelson et al. and those from the current analysis, restricted to SNP associations expected to appear in both datasets (pre-May 21, 2013 GWAS Catalog associations). Overlap is the number of items in common.

Table S15 presents data on Nelson et al. counts of distinct genes, trait MeSH headings, and SNPs compared to our results. The low overlap is striking. There are many sources of variation in the procedure.

1. Differences in the set of trait-SNP links.
2. Differences in LD expansion.
3. Differences in assigning LD SNPs to genes.
4. Differences in mapping traits to MeSH terms.

**Trait SNP links** As Nelson et al's table was constructed using the May 21, 2013 version of the GWASdb (including GWAS Catalog associations), we might expect their reported GWAS hits to be a subset of ours. Discrepancies between the two data sources are reduced but still substantial when comparing 2017 GWAS Catalog associations reported before May 21, 2013 and Nelson et al. associations with source "GWAS:A" (Figure S10, Table S16). Though some of this lack of overlap might be explained by differences in which SNPs could be mapped to genes or which studies were included, only 63% could be found in the current verison of the GWAS Catalog. 7% of studies account for these missing SNPs and tend to have larger total reported associations.

**LD expansion** Both LD expansions were conducted from EUR 1000 genomes [5], but Nelson et al. used pilot phase genomes and we used Phase 3. For associated SNPs common to both analyses, 1.3% of Nelson et al. LD SNPs were not recovered in the reanalysis. Nelson et al. do not provide an exhaustive enumeration of LD SNP-SNP links so the reverse calculation is not possible.

**LD SNP to Genes** Among LD SNPs in common between our dataset and the Nelson et al. dataset, 81% of LD SNP-Gene links appearing in the Nelson et al dataset also appear in our dataset. Previously-reported associations with eQTL evidence are less likely to be replicated in the current analysis (Table S17). Current associations based on eQTL evidence alone are also very unlikely to appear in the previous analysis (Table S18). Many SNPs with GTEx eQTL are not recognized by `eqtl.uchicago.edu`. Additionally, this is an older database and eQTLs may have been measured in fewer tissues and in studies with smaller sample sizes.

**Disease to MeSH Discrepancies** Nelson et al. provide mapping of many GWAS traits to MeSH terms in a supplementary table. From this table, around half of GWAS Catalog entries were mapped to a different disease term. It appears [20] mapped the `DISEASE.TRAIT` field rather than the `MAPPED_TRAIT` field as we did, and that they had a preference for mapping to disease terms even when the reported trait was not a disease (e.g. pulmonary function measurement mapped to lung diseases).

| eQTL | Percent in Current Table |
|---|---|
| FALSE | 85 |
| TRUE | 57 |

Table S17: Percent of LD SNP-Gene associations reported by Nelson et al. also found in the reanalysis by whether the association was an eQTL. Conditional on LD SNP being presence in both analyses.

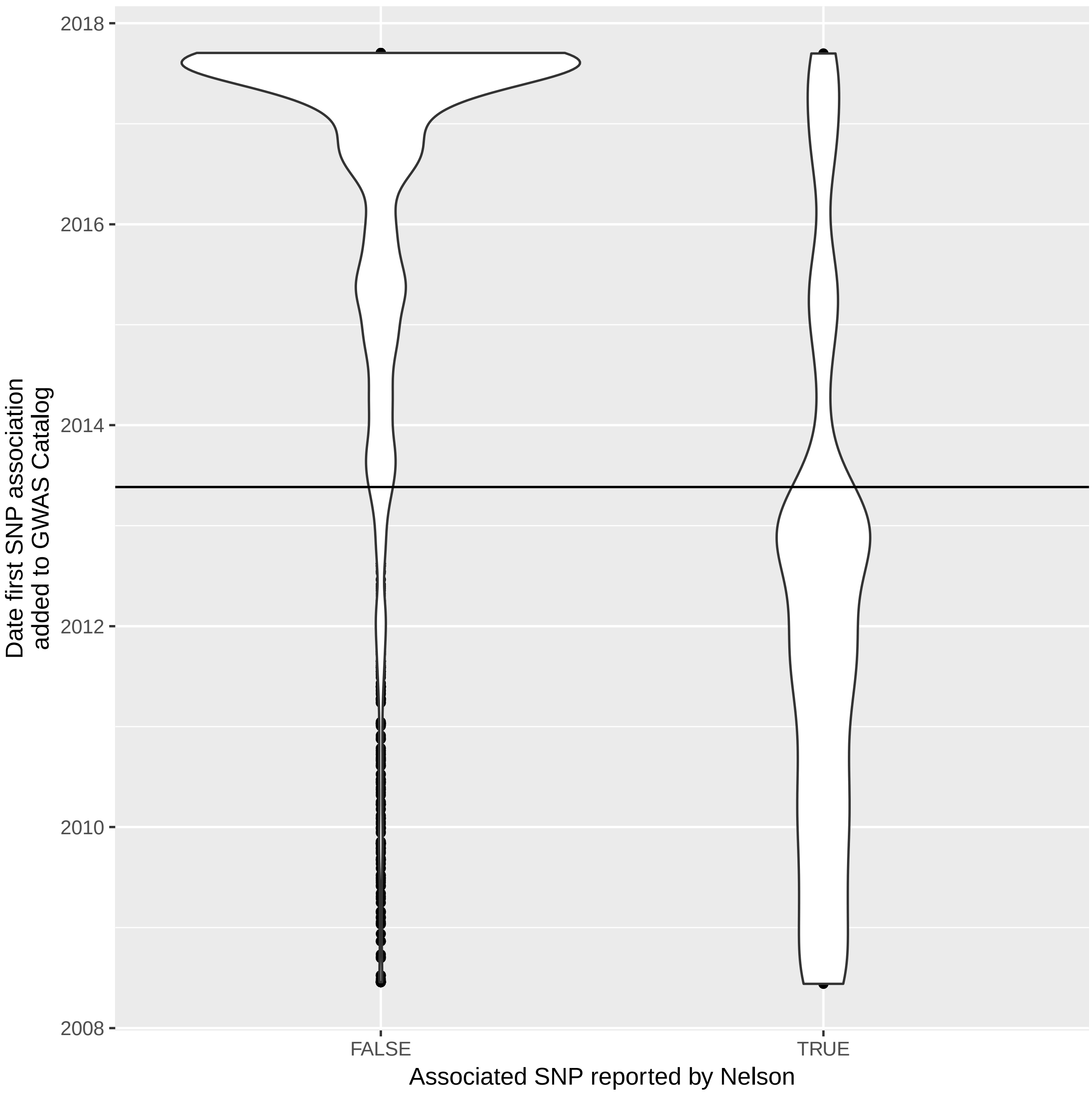

Figure S10: Date SNP associations appearing in the reanalysis were added to the GWAS Catalog by whether or not Nelson reported the association. Line shows the date of the GWASdb version used in [20].

| DHS | eQTL | distance | Percent in MN Table |
|---|---|---|---|
| TRUE | TRUE | TRUE | 83 |
| FALSE | TRUE | TRUE | 79 |
| TRUE | FALSE | TRUE | 69 |
| FALSE | FALSE | TRUE | 82 |
| TRUE | TRUE | FALSE | 0 |
| FALSE | TRUE | FALSE | 5 |
| TRUE | FALSE | FALSE | 9 |

Table S18: Percent of LD SNP-Gene associations found in reanalysis also reported by Nelson et al. subdivided by what evidence source(s) were used to link the LD SNP and gene. Conditional on LD SNP being presence in both analyses.

#### S3.2 Updating OMIM Dataset

**Inclusion Criteria** All phenotype associations with ensembl genes were obtained from the file `genemap2.txt` obtained from OMIM [19] (6275 total associations). Associations labelled as provisional were excluded (leaving 5800 associations). Drug response phenotypes and somatic mutations were also excluded (leaving 5522). Excluding drug response phenotypes is consistent with our GWAS protocol. Excluding somatic mutations was done to ensure tumor somatic mutations were not included, as whether cancer somatic variants predict successful drug targets is outside the scope of this analysis. Somatic mutations were recognized by a phenotype or mode of inheritance containing the word "somatic."

**Mapping to MeSH** OMIM phenotypes were mapped to MeSH using a hybrid automatic and manual assignment procedure. Automatic maps were generated by string matches to MeSH headings and supplementary concepts and by the Oxo tool [8] and reviewed for precision and accuracy. 92% of terms were automatically mapped and 8% of these were rejected after review. The remainder were manually mapped.

#### S3.3 Supplementary Results Using Updated Pipeline and Genetic Data

Figures S11 and S12 and Table S19 recreate Nelson et al. 2015 figures using updated genetic association and Pharmaprojects datasets. Enrichment of GWAS Catalog associated gene targets among approved drugs remains, but effect sizes are somewhat lower than originally reported (Figure S11). The pattern of increasing proportion of drug targets genetically associated with a phenotype similar to their indication through successive phases of clinical development is weaker for GWAS Catalog associations, though approved gene target-indication pairs still are the most likely to have genetic support for their targets (Figure S12). Table S19 replicates Table 1N (discussed in main text).

| | GWAS Catalog & OMIM | GWAS Catalog | OMIM |
|---|---|---|---|
| Preclinical to Phase I | 1 (1-1.1) | 1.1 (1-1.1) | 1 (1-1.1) |
| Phase I to Phase II | 1.1 (1.1-1.2) | 1.1 (1-1.1) | 1.2 (1.1-1.3) |
| Phase II to Phase III | 1.4 (1.3-1.5) | 1.1 (0.9-1.3) | 1.7 (1.5-1.8) |
| Phase III to Approved | 1.3 (1.2-1.4) | 1.1 (1-1.3) | 1.4 (1.3-1.5) |
| Phase I to Phase III | 1.6 (1.4-1.8) | 1.2 (1-1.4) | 2 (1.8-2.2) |
| Phase I to Approved | 2.1 (1.8-2.4) | 1.4 (1.1-1.7) | 2.7 (2.4-3.1) |

Table S19: Replication of Table 1N (association between genetic evidence and historical progression) from updated GWAS Catalog and OMIM genetic association dataset and updated pipeline data (Full Data). Risk ratio p(approved | genetic support)/p(approved | no genetic support) and bootstrap 95% confidence intervals.

**eQTL p-value cutoff** The eQTL p-value raw cutoff was chosen somewhat arbitrarily as $10^{-6}$ in any GTEx tissue [11]. Given the number of comparisons made, this cutoff is not very stringent and could possibly weaken the estimated effect of GWAS genetic evidence by adding noise to the list of gene-trait links. Table S20 estimates of the effect of GWAS genetic evidence in the Full Data set using only eQTL with $p$-value less than $10^{-12}$. Using this more stringent cutoff reduces the number of gene-trait links from 28366 to 24681.

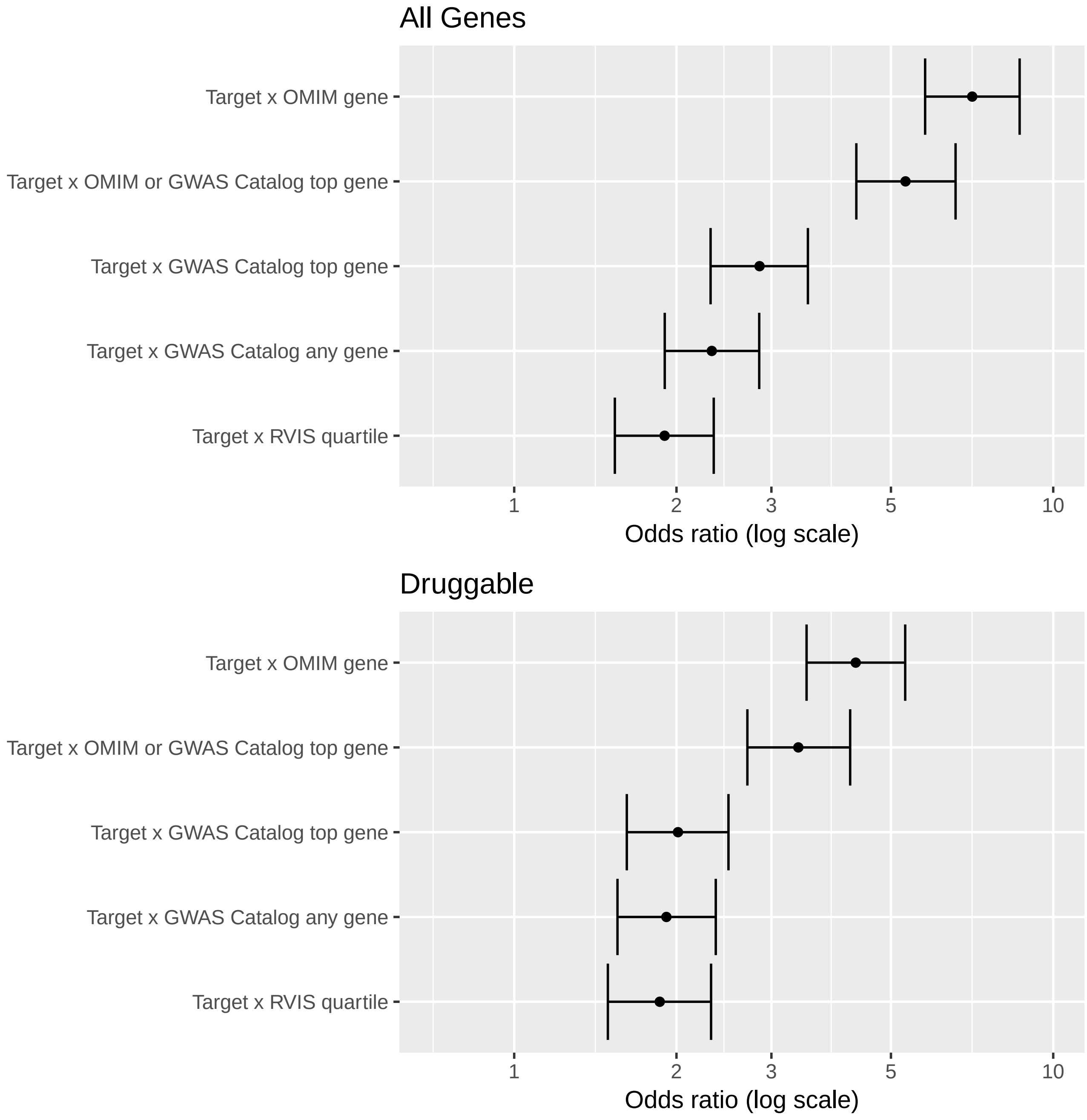

Figure S11: Replication of Figure 2N from updated GWAS Catalog genetic association dataset and updated pipeline data. Figure shows the enrichment of approved drug targets among genes with human genetic associations.

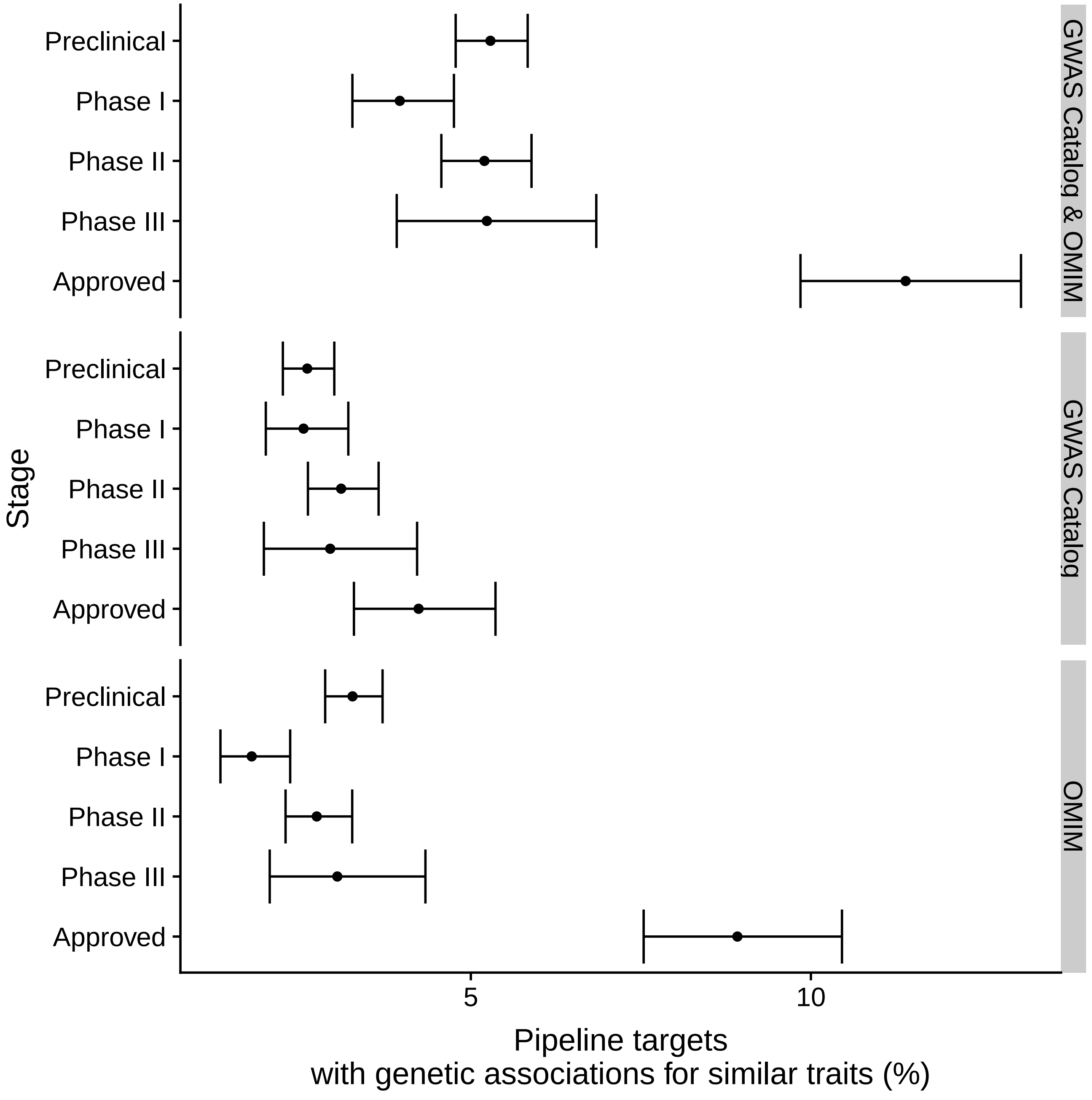

Figure S12: Replication of Figure 3Nb from updated GWAS Catalog genetic association dataset and updated pipeline data. Figure shows the proportion of gene target-indication pairs with genetic associations for similar traits by pipeline phase and association source.

| | GWAS Catalog & OMIM | GWAS Catalog | OMIM |
|---|---|---|---|
| Preclinical to Phase I | 1 (0.9-1.1) | 1 (1-1.1) | 1 (0.9-1) |
| Phase I to Phase II | 1.1 (1.1-1.2) | 1.1 (1-1.1) | 1.2 (1.1-1.2) |
| Phase II to Phase III | 1.5 (1.3-1.6) | 1.2 (1-1.4) | 1.7 (1.5-1.9) |
| Phase III to Approved | 1.3 (1.2-1.4) | 1.2 (1-1.3) | 1.4 (1.3-1.5) |
| Phase I to Phase III | 1.6 (1.5-1.8) | 1.2 (1-1.4) | 2 (1.8-2.3) |
| Phase I to Approved | 2.2 (1.9-2.4) | 1.4 (1.1-1.7) | 2.8 (2.4-3.2) |

Table S20: Replication of Nelson et al. Table 1 from updated GWAS Catalog and OMIM genetic association dataset and updated pipeline data, eQTL p-value cutoff $10^{-12}$.

### S4 Trait-Indication Similarity: Supplementary Methods And Results

#### S4.1 Methods

##### S4.1.1 Information Content

[21] provide an overview of semantic similarity measures in biomedical ontologies. Resnik and Lin similarities are both information content (IC) based similarity measures. IC can be computed externally from term frequencies in a corpus or through the structure of the ontology itself [25]. We computed information content using the `descendants_IC` function from the `R` package `ontologySimilarity` [9]. Let $c$ be a term. `descendants_IC` computes information content as

$$IC(c) = -\log\left(\frac{a(c)}{N}\right)$$

Where $N$ is the total number of terms in the ontology and $a(c)$ is the number of terms for which $c$ is an ancestor (including itself). We chose this approach over corpus-based similarity available from `UMLS::Similarity` used by Nelson et al because the latter is not able to compute information content for terms that do not appear in 2009 MeSH, and we mapped to 2017 MeSH. Additionally, we found that corpus-based MeSH heading similarities computing using information content from the most current version of PubMed deviated more from Nelson et al. original values than did our values computed using descendant IC for similarities near the 0.7 cutoff used in the original analysis (not shown).

##### S4.1.2 Similarity

Resnik similarity between terms $c_1$ and $c_2$ is computed as

$$\text{sim}_{\text{res}}(c_1, c_2) = IC(MICA(c_1, c_2)) \tag{1}$$

where $MICA(c_1, c_2)$ is the common ancestor of $c_1$ and $c_2$ with the highest information content.

Lin similarity is computed as

$$\text{sim}_{\text{lin}}(c_1, c_2) = \frac{2 \times IC(MICA(c_1, c_2))}{IC(c_1) + IC(c_2)} \tag{2}$$

Nelson et al. quantified semantic similarity as the average of Resnik [24] and Lin [15] similarities each rescaled to between zero and one. Lin similarity is between zero and one, and does not need to be rescaled. We computed rescaled Resnik similarity using the following formula.

$$\text{sim}_{\text{res,norm}}(c_1, c_2) = \begin{cases} \frac{IC(MICA(c_1,c_2))}{\max_{c \in \mathcal{C}} IC(c)} & c_1 \neq c_2 \\ 1 & c_1 = c_2 \end{cases} \tag{3}$$

Figure S13 shows a portion of the MeSH vocabulary for heart diseases, with parent term Cardiovascular Diseases.

Table S21 shows similarities computed between trait pairs in Figure S13. For example, we compute similarities between Cardiomyopathies and Arrhythmias, Cardiac as follows.

$$\begin{aligned} \text{sim}_{\text{res,norm}}(\text{Cardiomyopathies, Arrhythmias, Cardiac}) &= \frac{IC(\text{Heart Diseases})}{\max_{c \in \mathcal{C}} IC(c)} \\ &= \frac{5.07}{10.26} \\ &= 0.49 \end{aligned}$$

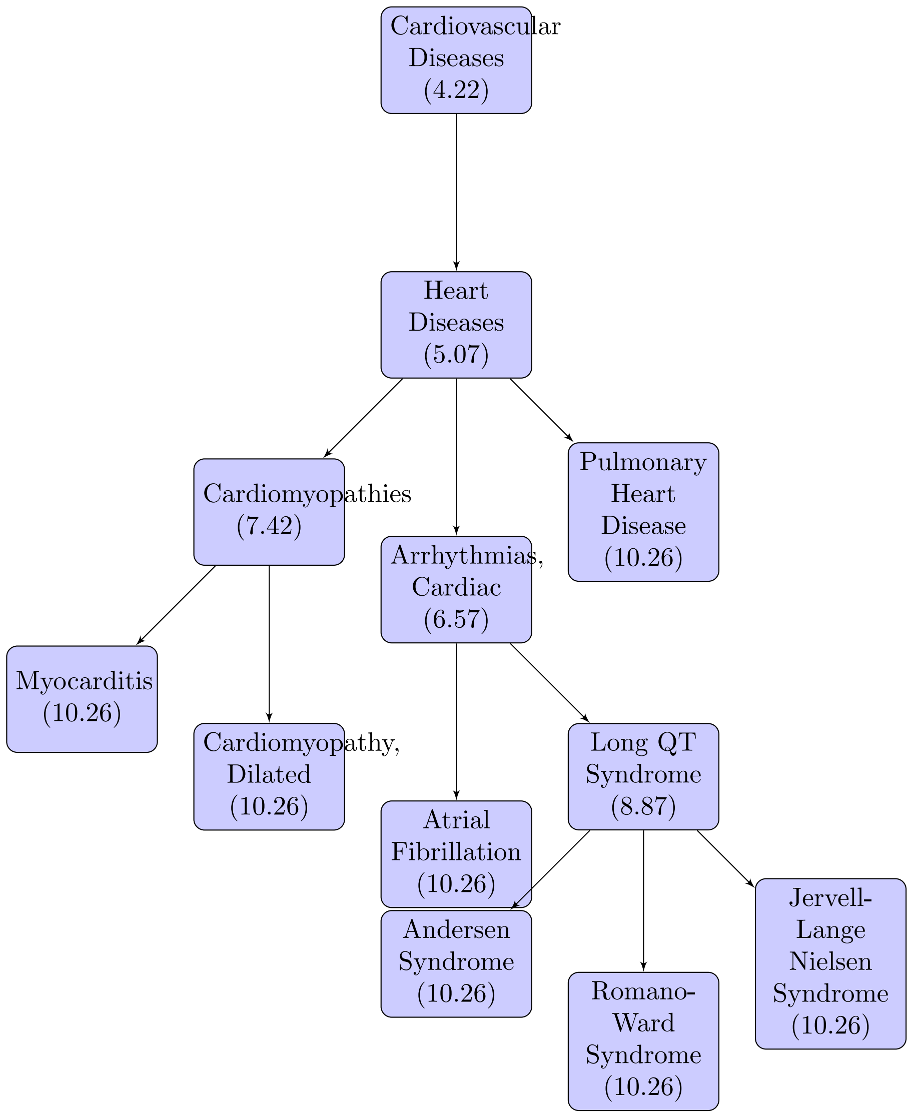

Figure S13: A portion of the MeSH vocabulary used to illustrate semantic similarity. Information contents from the number of descendants (computed from the entire ontology, not just the portion shown here) are given in parentheses.

| Term 1 | Term 2 | $\text{sim}_{\text{res}}$ | $\text{sim}_{\text{lin}}$ | $\text{sim}_{\text{res,norm}}$ | $\text{sim}_{\text{avg}}$ |
|---|---|---|---|---|---|
| Heart Diseases | Heart Diseases | 5.07 | 1 | 1 | 1 |
| Andersen Syndrome | Andersen Syndrome | 10.26 | 1 | 1 | 1 |
| Cardiomyopathies | Arrhythmias, Cardiac | 5.07 | 0.73 | 0.49 | 0.61 |
| Cardiomyopathy, Dilated | Romano-Ward Syndrome | 5.07 | 0.49 | 0.49 | 0.49 |
| Andersen Syndrome | Romano-Ward Syndrome | 8.87 | 0.86 | 0.86 | 0.86 |
| Arrhythmias, Cardiac | Atrial Fibrillation | 6.57 | 0.78 | 0.64 | 0.71 |

Table S21: Similarity between different pairs of traits computed using Resnik similarities, Lin similarities, normalized Resnik similarities, and average of normalized Resnik and Lin similarities using number of descendants to compute information content.

$$
\begin{aligned}
\text{sim}_{\text{lin}}(\text{Cardiomyopathies, Arrhythmias, Cardiac}) &= \frac{2 \times IC(\text{Heart Diseases})}{IC(\text{Cardiomyopathies}) + IC(\text{Arrhythmias, Cardiac})} \\
&= \frac{2 \times 5.07}{7.42 + 6.57} \\
&= 0.71
\end{aligned}
$$

##### S4.2 Comparing Nelson et al MeSH similarities with this study

Figure S14 shows the correlation between average similarity as reported by Nelson et al. and our calculated values. Note that a similarity cutoff of 0.7 in the Nelson et al. analysis (red vertical line) is equivalent to a somewhat higher cutoff in our analysis. Using linear regression, we estimate the similarity cutoff of 0.7 used in the original analysis is equivalent to a similarity cutoff of 0.73 in our work, motivating our choice of cutoff. Clinical progression probability estimates using the previous cutoff value of 0.7 (Figure S15) are similar to those reported in the main text.

##### S4.3 Effect of manually assigned similarity on approval

The MeSH vocabulary rarely provides links between diseases and related quantitative traits. To be able to use quantitative trait association studies as supporting genetic evidence, Nelson et al. manually assigned similarities to 320 trait-indication pairs. To assess the effect of manually assigned similarities on estimates of the effect of genetic evidence, we used supplementary tables to recreate the MeSH similarity matrix without manually assigned similarities, and recomputed risk ratios for the effect of GWAS genetic evidence on gene target-indication pair progression from Phase I to approval. We also did the same for our updated datasets. Finally, we looked at the effect of using only manually assigned similarities (setting all other similarities equal to zero) on our estimates. For consistency, we used the same set of well-studied MeSH indications in all estimates (rather than recomputing them for each similarity matrix). OMIM associations were not supported by genetic evidence based on manually assigned similarities, so there is no effect on estimates of OMIM genetic evidence, and this is not shown.

Results are shown in Figure S16. Risk ratios of progression from Phase I to approval for gene target-indication pairs supported by manually assigned traits are very high at all similarity cutoff values (chosen as 0.5, 0.75, and 0.9 as these were the three manually assigned values). These appear to be driving most of the observed significant, positive effect of GWAS genetic evidence, though we still see some non-significant, positive odds ratios at higher similarity cutoff values. One hypothesis for this effect is that manually assigned similarities occur for a non-random set of indications, and these indications are more likely to be successful. However, we do not find differences in approval rate for these indications. It is also possible that automatically assigned MeSH similarities are more reliable indications of a genuine biological link between traits than traits with comparable automatically assigned similarities because of the expert knowledge going into them or that quantitative trait associations are more informative about disease mechanisms. However, the predictive power of manually assigned similarities does not replicate well using new GWAS Catalog associations to predict success (analysis New Genetic in Figure S16), suggesting use of more objective methods is advisable.

#### S5 Modeling Drug Success Probability: Supplementary Methods And Results

In preliminary modeling work, we found GWAS genetic evidence *decreased* approval probability when the associated trait was dissimilar to the drug indication. In contrast, gene target-indication pairs with OMIM genetic evidence were significantly more likely to have an approved drug even if the reported trait was dissimilar to the drug indication. We suspected differences between targets linked to GWAS Catalog traits, targets linked to OMIM traits, and targets found in neither source could explain this. In this section, we fit logistic regression models to gene target-indication pair-level approval data with predictors whether or not a target is a GWAS or an OMIM gene, maximum similarity of a trait associated with the target to the indication, and target and indication-level properties (see Methods).

##### S5.1 Predictors

**Highest level MeSH** All Pharmaprojects indications are associated with one or more top level terms in the MeSH disease or psychiatry and psychology hierarchy (for example, Neoplasms and Immune System Diseases). Previous

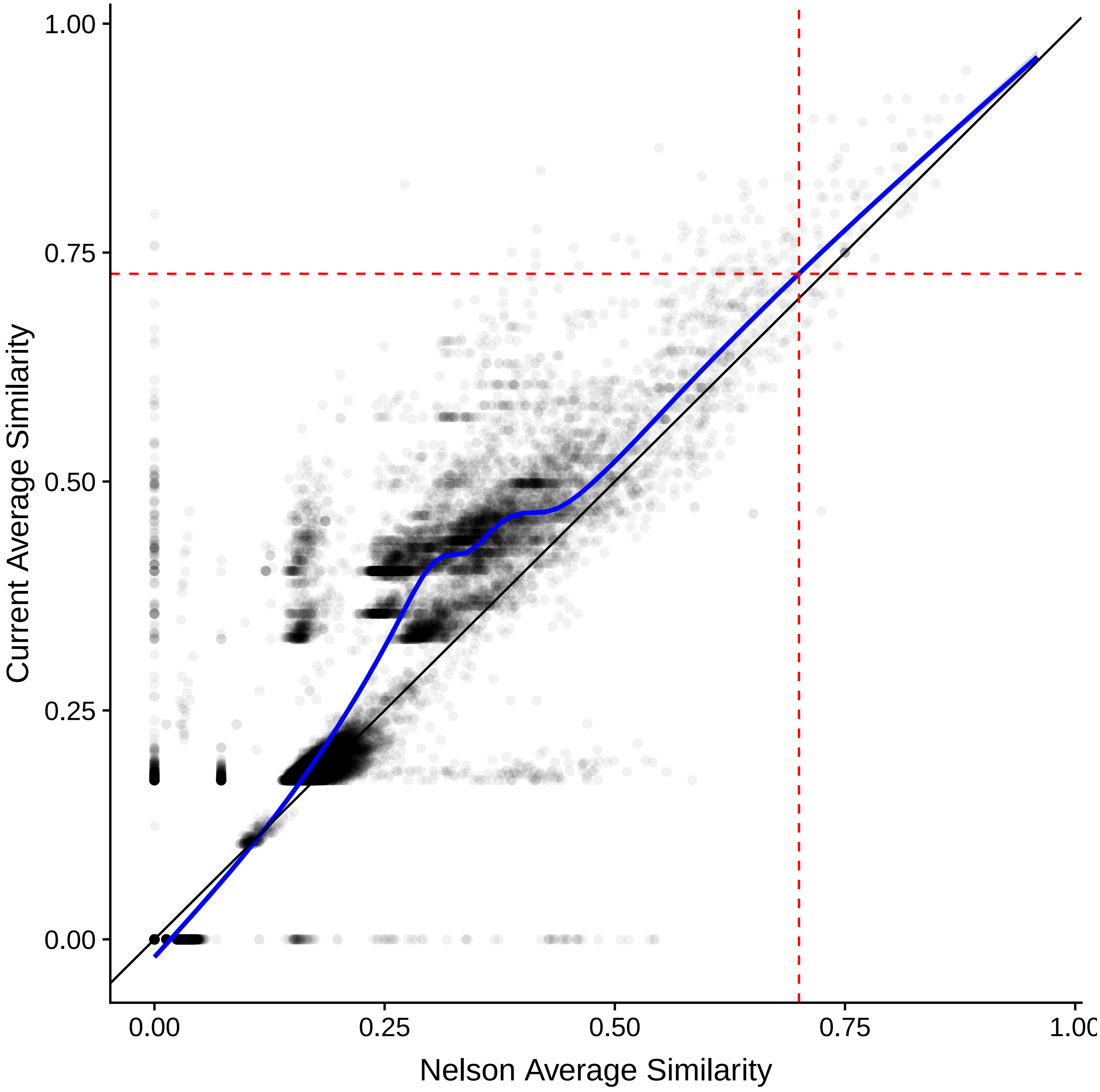

Figure S14: Nelson et al. average similarity versus average similarity computed in this analysis. Black points show a random sample of 50,000 trait pairs for which both similarities were available. Blue line shows smoothed relationship estimated using `gam` using all possible similarity pairs. Dashed red lines show old and new cutoff values.

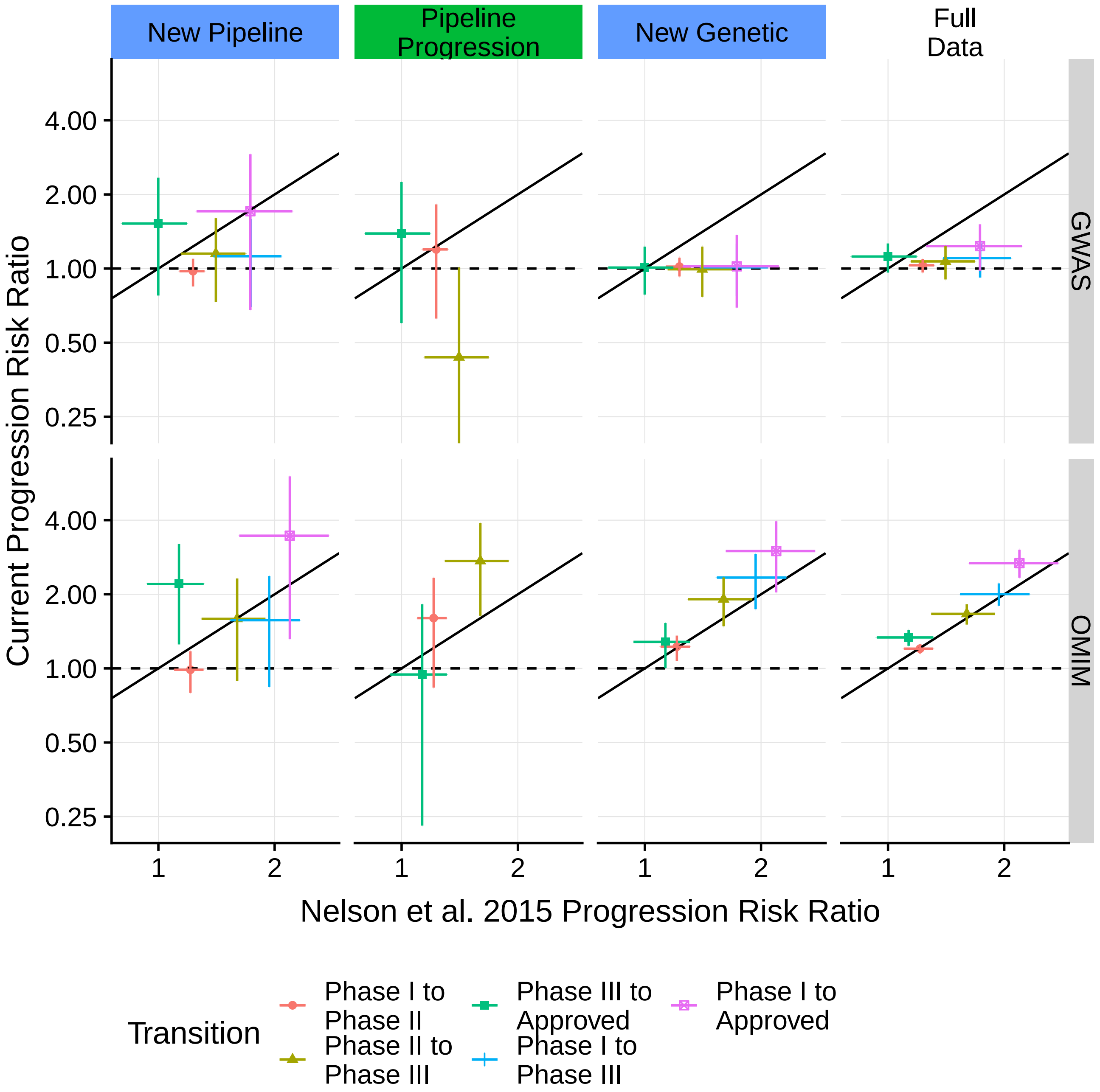

Figure S15: Estimated effect of genetic evidence on pipeline progression. Main text Figure 1b computed with similarity cutoff 0.7. This cutoff was originally used in Table 1N, but our main text figure uses similarity cutoff 0.73 because of systematic differences in computed similarities.

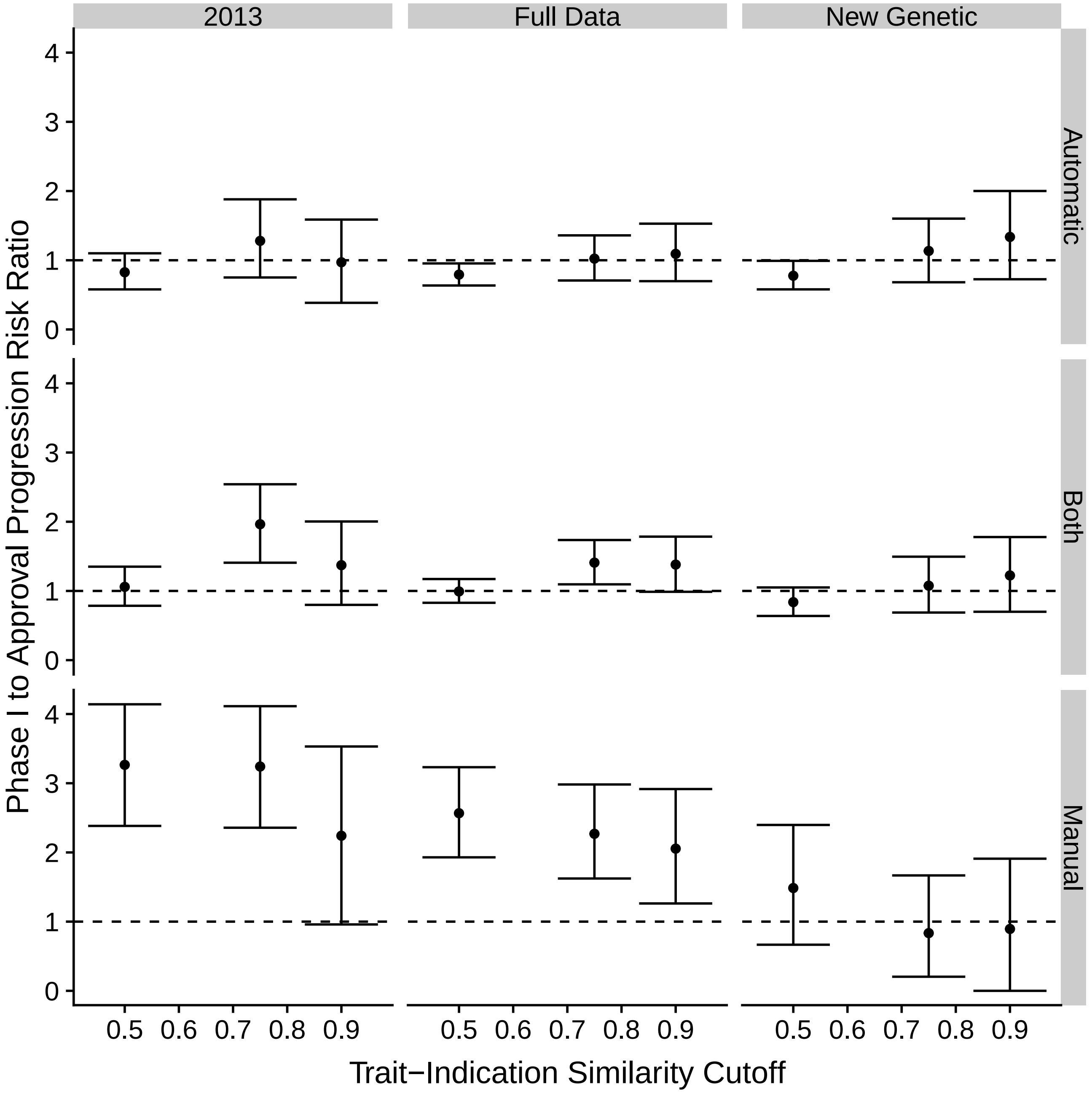

Figure S16: Risk ratio of Phase I to approval progression for gene target-indication pairs with and without genetic evidence for different values of the MeSH similarity cutoff split by whether MeSH similarity is automatically assigned, manually assigned, or using both automatic and manually assigned similarities (default). Full Data and New Genetic are as described in main text. 2013: computed from supplementary tables.

work has shown differences in approval probability for drugs by indication category, for example lower approval probabilities for oncology indications [12]. Figure S17 shows approval probability against GWAS target probability for different top-level MeSH terms. Neoplasms, the most common top-level MeSH in the dataset, is associated with both lower approval and higher probability of a GWAS associated target. We use presence or absence of each top level MeSH term annotation to predict gene target-indication pair success, so gene target-indication pairs may have more than one top-level MeSH term.

**GO Annotation** Previous work has shown differences in success probability among target classes (e.g. kinases and GPCR) [27]. Pharmaprojects provides functional annotations which seemed to be predictive of success, but were not available for all targets. We translated these annotations to Gene Ontology (GO) [2][6] terms when possible, found all descendants of these terms using the `goatools` Python module [13], and then annotated each gene target with a term if it was annotated with that term or a descendant term. GO term annotations were obtained using biomaRt [28]. Figure S18 shows variation in approval and GWAS links by target class.

**Variance Tolerance** Residual Variation Intolerance scores measure tolerance of functional variation [22]. Nelson et al. showed genes with lower values of this score are more likely to be the targets of new drugs, but that accounting for this has little effect on the estimated effect of genetic evidence. We obtained RVIS scores from the supplementary tables of [22], and confirm a weak negative association of RVIS percentile with approval (Figure S19).

**Time of First Attempt of Target** One hypothesis for lower success of GWAS gene targets is that GWAS is inspiring the choice of new targets that are either less druggable or have not had enough time to produce an approved drug. For most gene targets in the Pharmaprojects database, we can compute the earliest date at which a drug with that target was added to the database (an estimate of the length of time the target has been under development) and use this to predict whether that target occurs in least one successful drug. Figure S20 shows the relationship between the date the target was added and approval. Targets that have been under development for longer are more likely to have an approved drug. The complete lack of approvals for target added since 2013 likely reflects the timescale of clinical development.

#### S5.2 Supplementary Results

Figure S21 shows coefficient estimates (intercept, linear, and quadratic effects) for genetic evidence using different predictor subsets. The changes in coefficient estimates from including different covariates are meaningful, but the coefficient estimates are not comparable across GWAS and OMIM as trait similarity has been centered and scaled separately for each data source. Figure S22 shows the effect of including gene and indication-level predictors on the estimated relationship between indication-trait similarity and approval, which allows meaningful comparison between GWAS and OMIM. Without accounting for covariates, genes connected to a GWAS trait are systematically less likely to be the target of an approved drug relative to genes reported in OMIM when the trait and drug indication are highly dissimilar. Adding target and gene level properties to the model reduces this difference. In particular, genes reported in OMIM are more likely to be the targets of older drugs, which are more likely to be approved. Including target class annotations in the model has the largest effect on GWAS coefficient estimates. Adding all target and indication level predictors improves the WAIC model selection criterion over models without such predictors, or with only a single target and indication level predictor.

Figure S23 shows the effect of LD expansion threshold and trait similarity on the effect of a GWAS genetic association linked to a highly or moderately deleterious variant on gene target-indication pair approval. Results are not highly sensitive to the choice of threshold but in the direction expected: a genetic association for a similar trait has a larger effect on success if the deleterious SNP is more closely linked to the top GWAS variant.

In the previous section, we showed that manually assigned similarities are highly predictive of success for 2013 data, but this does not extend to our independent replication sets. We repeat our statistical analysis without manually assigned similarities to address our concerns about their predictive validity. We find a small reduction in estimated approval odds ratios for gene target-indication pairs with genetic evidence, but the relationship is not qualitatively different (Figure S24)

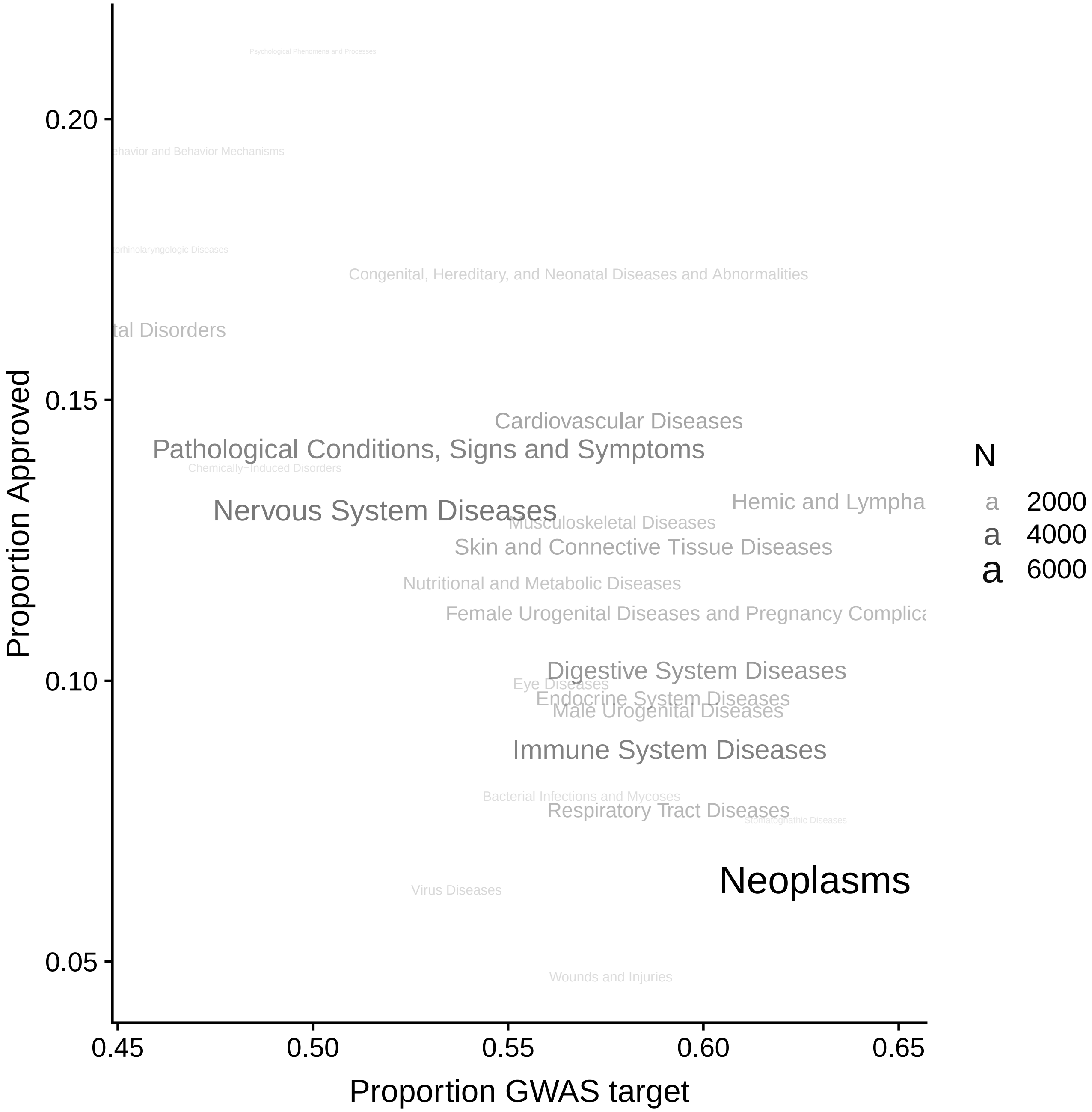

Figure S17: Proportion of gene target-indication pairs approved against proportion of gene target-indication pairs with a GWAS Catalog associated target by indication class. Larger text designates indication classes with more gene target-indication pairs. Only classes with 50 or more pairs are shown.

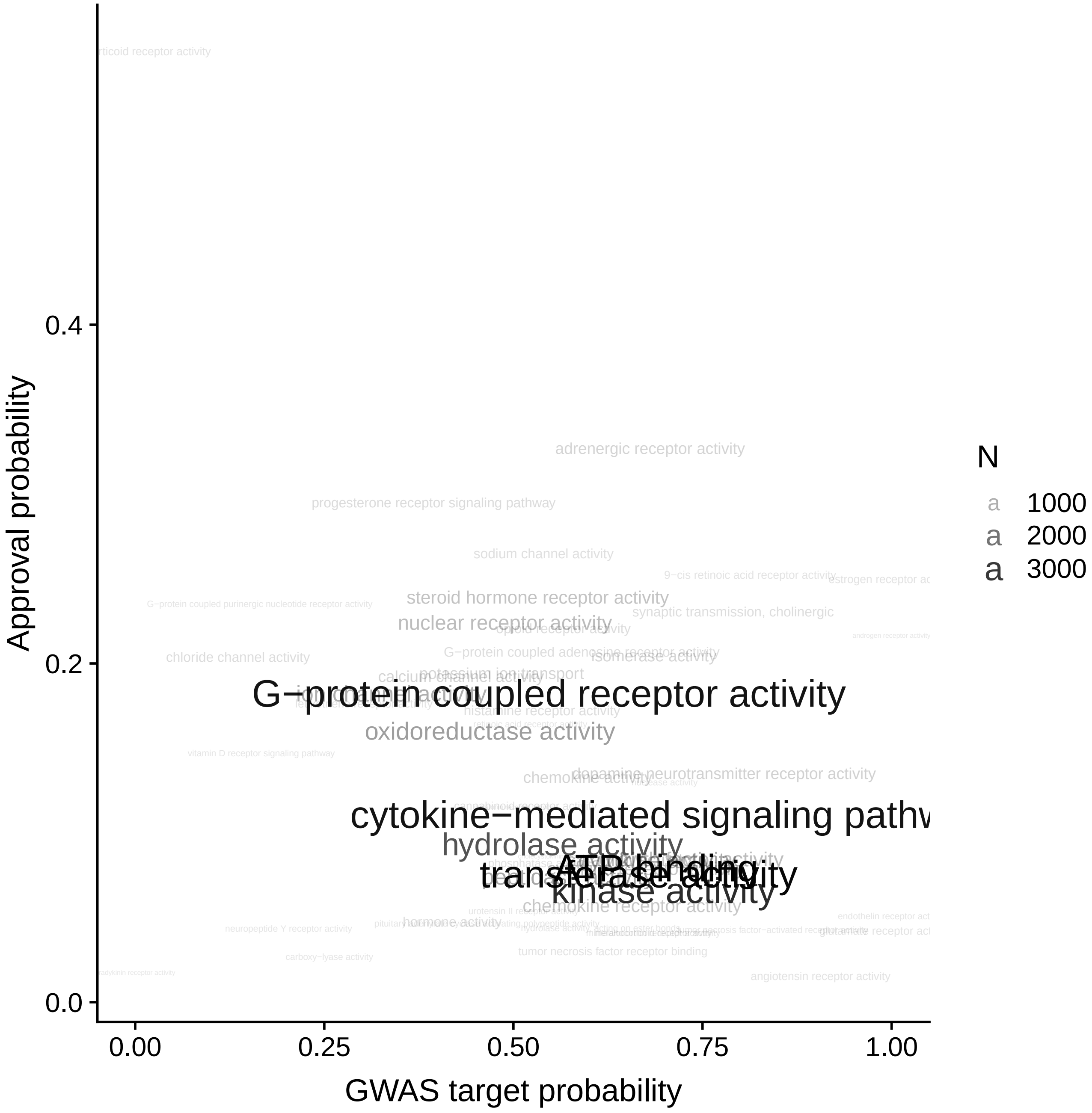

Figure S18: Proportion of gene target-indication pairs approved against proportion of gene target-indication pairs with a GWAS Catalog associated target by target class (GO terms). Larger text designates target classes with more gene target-indication pairs. Only classes with 50 or more pairs are shown.

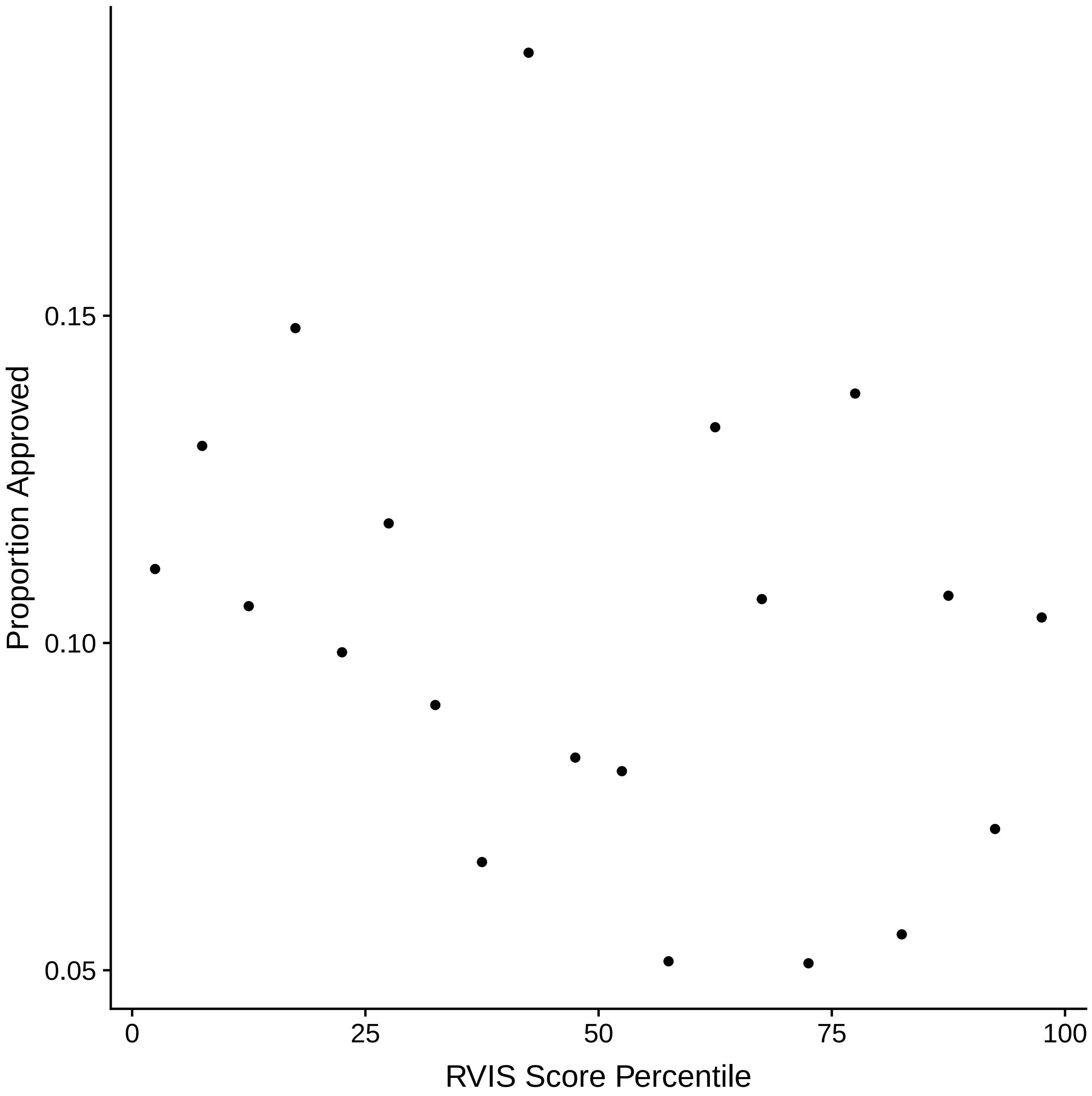

Figure S19: Proportion of approved gene target-indication pairs binned by target RVIS score percentile.

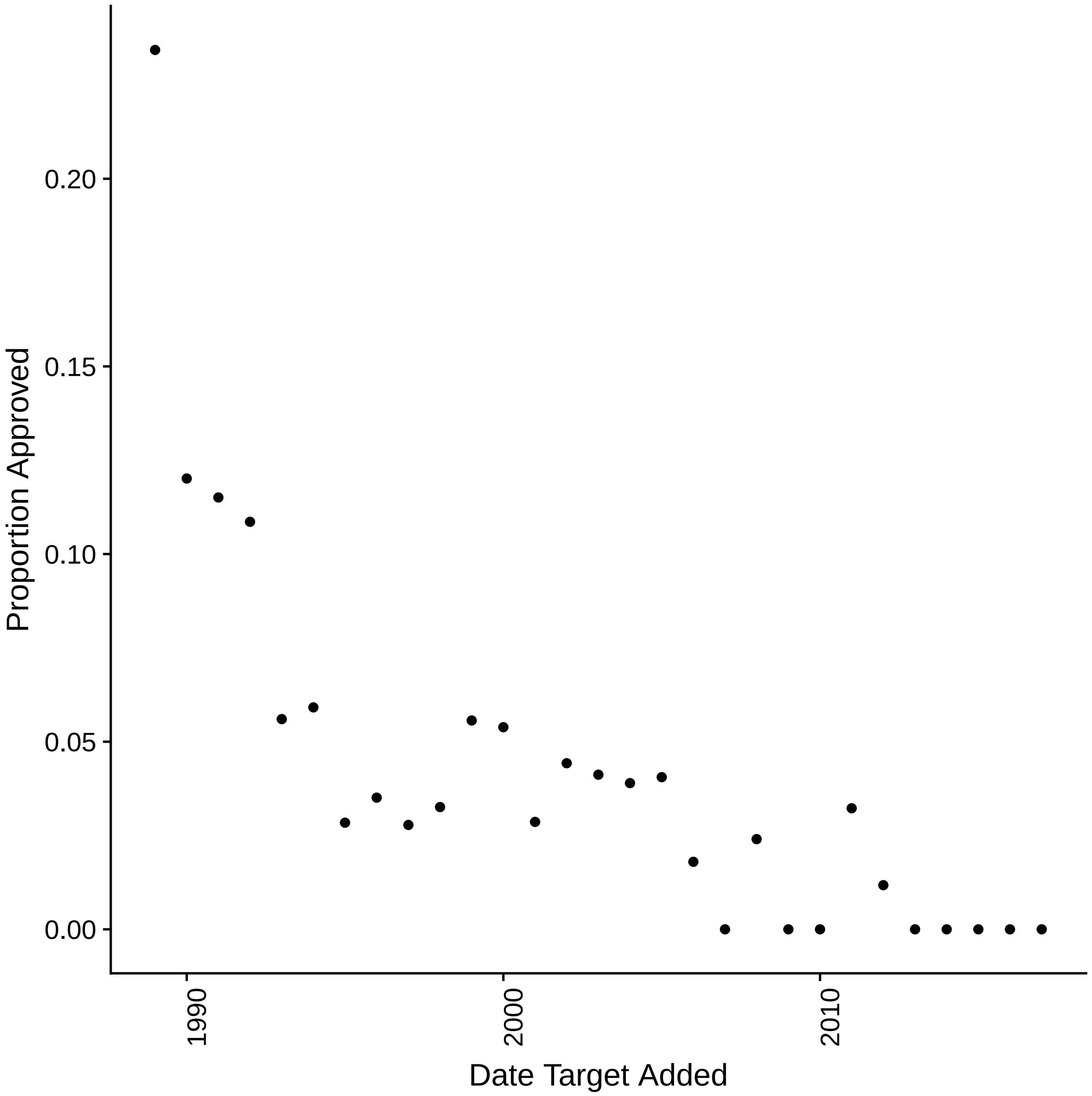

Figure S20: Proportion of approved gene target-indication pairs binned by date first drug with target added to Pharmaprojects.

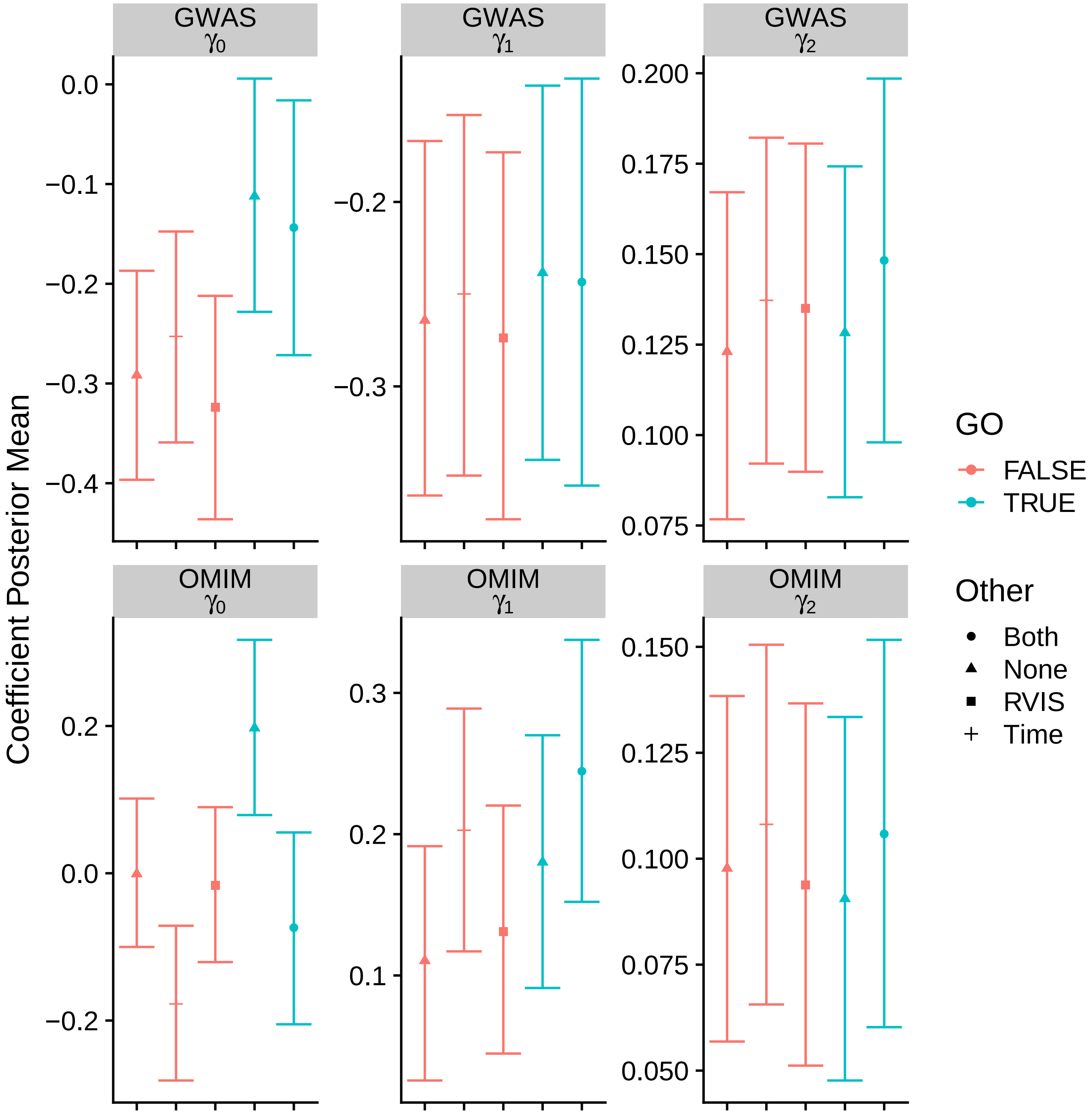

Figure S21: Coefficient estimates for the effect of genetically associated trait similarity on gene target-indication pair approval using different predictor subsets. See methods for details coefficient definitions. Note coefficients apply to centered and scaled trait-indication similarity so that the intercept for GWAS genetic evidence is the effect of genetic evidence at the mean value of GWAS trait similarity. GO = Gene Ontology terms, RVIS = RVIS score, Time = Time since target entered development.

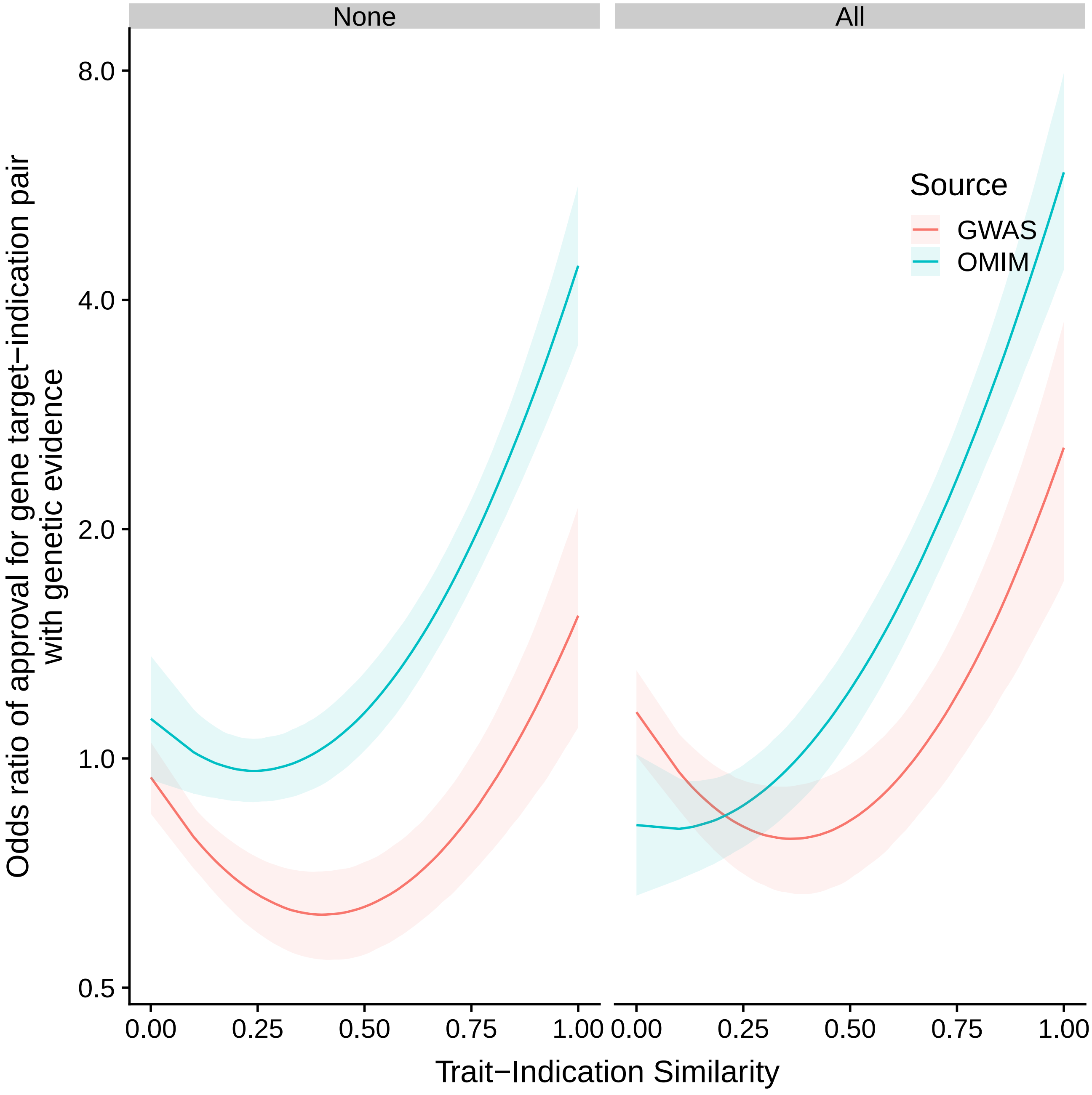

Figure S22: Effect of including predictors on the estimated relationship between indication-trait similarity and approval. Estimated odds ratio of gene target-indication pair attaining approval, as a function of similarity between drug indication and the most similar trait associated with the target. Posterior median and pointwise 95% credible interval from Bayesian logistic regression.

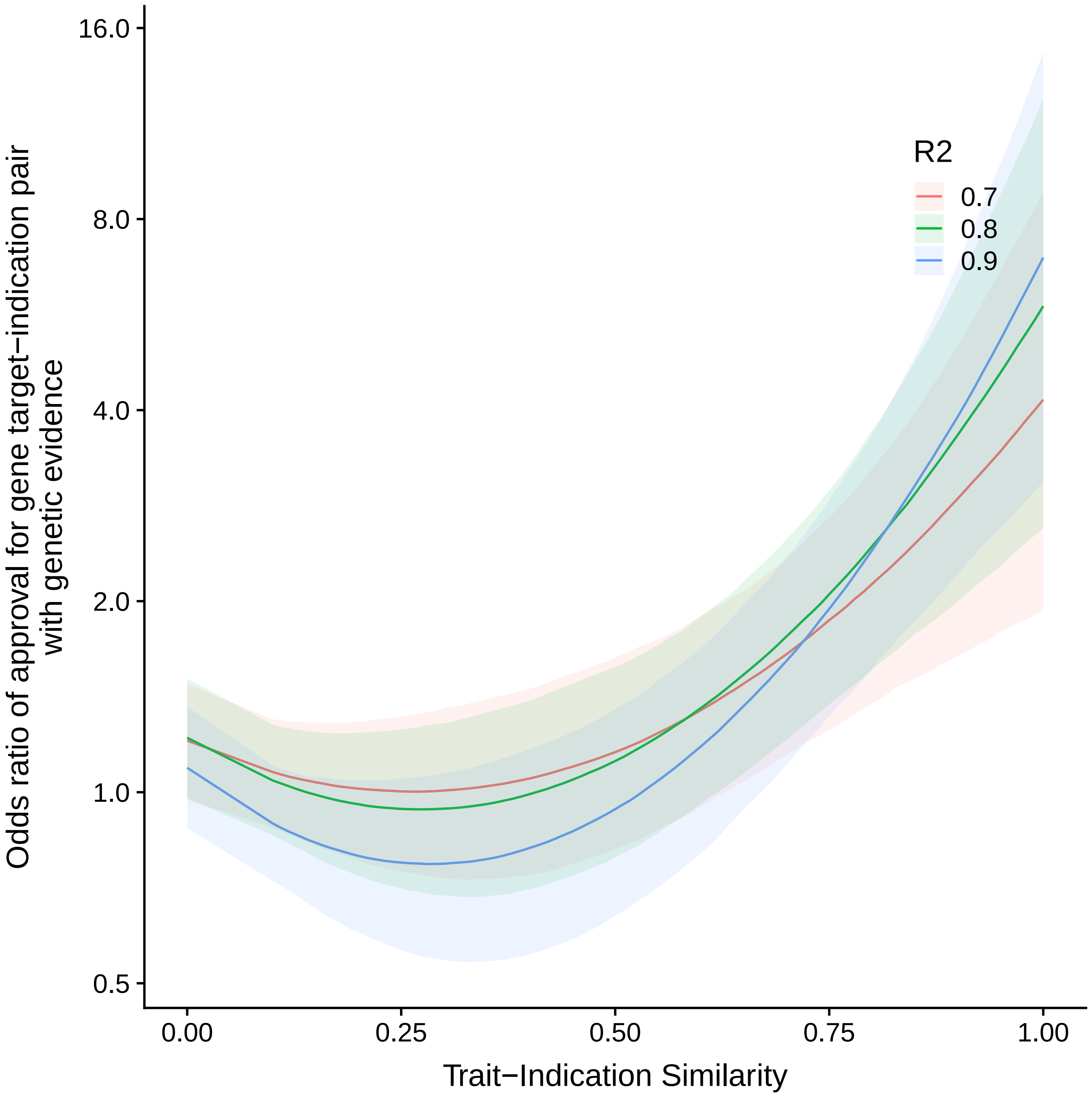

Figure S23: Effect of LD expansion threshold on the predicted success odds ratio of a drug gene target-indication pair supported by a GWAS high-moderate deleterious variant. Estimated odds ratio of gene target-indication pair attaining approval, as a function of similarity between drug indication and the most similar trait associated with the target. Colors show values for different values of the LD expansion threshold. Posterior median and pointwise 95% credible interval from Bayesian logistic regression.

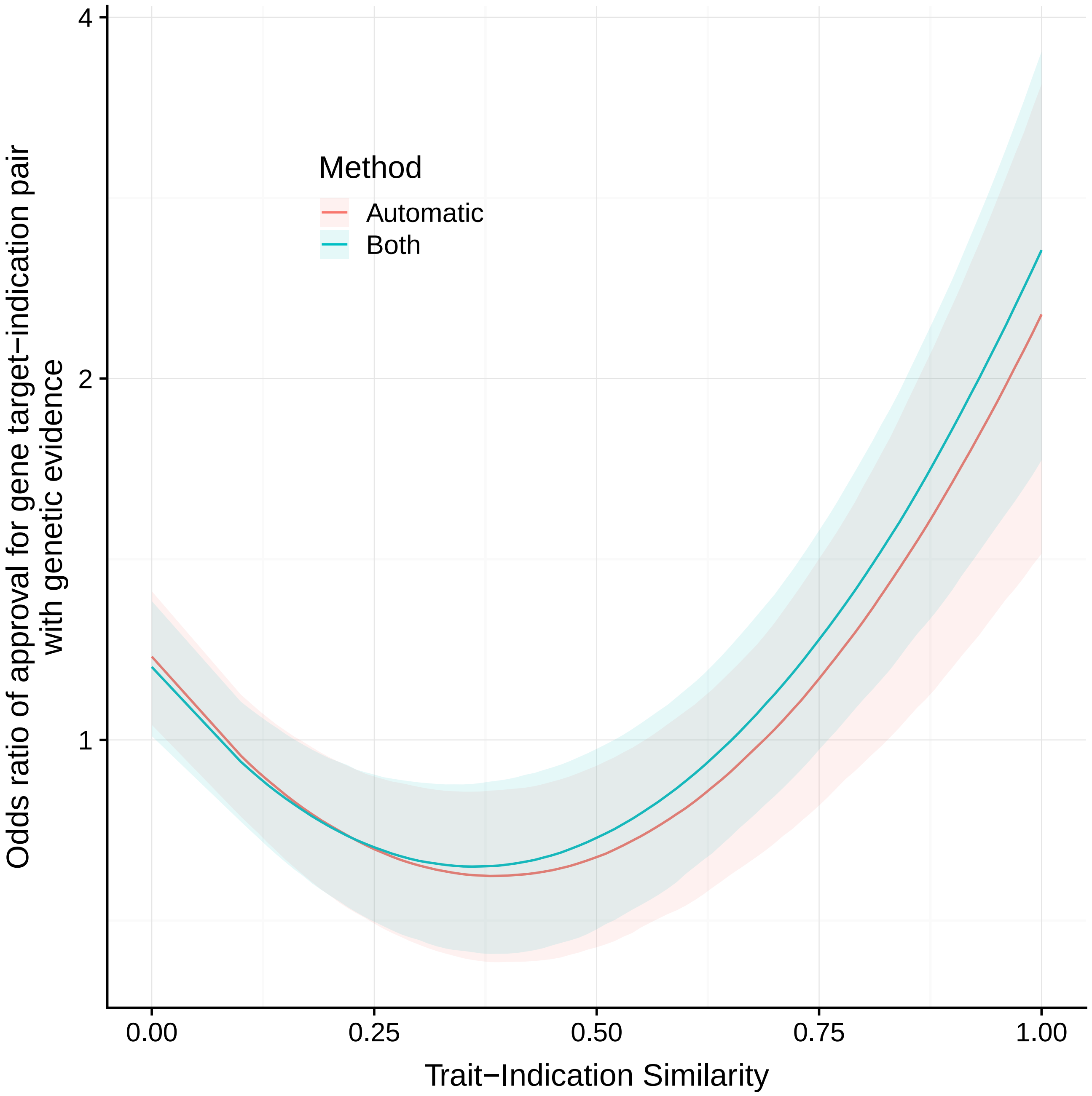

Figure S24: Effect of excluding manually assigned trait similarities on estimated relationship between GWAS genetic support and approval. Estimated odds ratio of gene target-indication pair attaining approval, as a function of similarity between drug indication and the most similar trait associated with the target. The two colors correspond to estimates when using and when excluding manually assigned similarities. Posterior median and pointwise 95% credible interval from Bayesian logistic regression.

[2] Michael Ashburner, Catherine A Ball, Judith A Blake, David Botstein, Heather Butler, J Michael Cherry, Allan P Davis, Kara Dolinski, Selina S Dwight, Janan T Eppig, et al. Gene Ontology: tool for the unification of biology. *Nature Genetics*, 25(1):25, 2000.

[3] Alan P Boyle, Eurie L Hong, Manoj Hariharan, Yong Cheng, Marc A Schaub, Maya Kasowski, Konrad J Karczewski, Julie Park, Benjamin C Hitz, Shuai Weng, et al. Annotation of functional variation in personal genomes using regulomedb. *Genome Research*, 22(9):1790–1797, 2012.

[4] P. Cingolani, A. Platts, M. Coon, T. Nguyen, L. Wang, S.J. Land, X. Lu, and D.M. Ruden. A program for annotating and predicting the effects of single nucleotide polymorphisms, SnpEff: SNPs in the genome of Drosophila melanogaster strain w1118; iso-2; iso-3. *Fly*, 6(2):80–92, 2012.

[5] 1000 Genomes Project Consortium et al. A global reference for human genetic variation. *Nature*, 526(7571):68–74, 2015.

[6] Gene Ontology Consortium. Expansion of the Gene Ontology knowledgebase and resources. *Nucleic Acids Research*, 45(D1):D331–D338, 2016.

[14] Mulin Jun Li, Panwen Wang, Xiaorong Liu, Ee Lyn Lim, Zhangyong Wang, Meredith Yeager, Maria P Wong, Pak Chung Sham, Stephen J Chanock, and Junwen Wang. GWASdb: a database for human genetic variants identified by genome-wide association studies. *Nucleic Acids Research*, 40(D1):D1047–D1054, 2011.

[25] Nuno Seco, Tony Veale, and Jer Hayes. An intrinsic information content metric for semantic similarity in wordnet. In *ECAI*, volume 16, page 1089, 2004.

[26] Nathan C Sheffield, Robert E Thurman, Lingyun Song, Alexias Safi, John A Stamatoyannopoulos, Boris Lenhard, Gregory E Crawford, and Terrence S Furey. Patterns of regulatory activity across diverse human cell types predict tissue identity, transcription factor binding, and long-range interactions. *Genome Research*, 23(5):777–788, 2013.

[27] Hsin-Pei Shih, Xiaodan Zhang, and Alex M Aronov. Drug discovery effectiveness from the standpoint of therapeutic mechanisms and indications. *Nature Reviews Drug Discovery*, 17(1):19, 2018.

[28] Damian Smedley, Syed Haider, Steffen Durinck, Luca Pandini, Paolo Provero, James Allen, Olivier Arnaiz, Mohammad Hamza Awedh, Richard Baldock, Giulia Barbiera, et al. The BioMart community portal: an innovative alternative to large, centralized data repositories. *Nucleic Acids Research*, 43(W1):W589–W598, 2015.

[29] Alex H Wagner, Adam C Coffman, Benjamin J Ainscough, Nicholas C Spies, Zachary L Skidmore, Katie M Campbell, Kilannin Krysiak, Deng Pan, Joshua F McMichael, James M Eldred, et al. DGIdb 2.0: mining clinically relevant drug–gene interactions. *Nucleic Acids Research*, 44(D1):D1036–D1044, 2015.
